# An evolutionarily diverged mitochondrial protein controls biofilm growth and virulence in *Candida albicans*

**DOI:** 10.1101/2020.09.29.318048

**Authors:** Zeinab Mamouei, Shakti Singh, Bernard Lemire, Yiyou Gu, Abdullah Alqarihi, Sunna Nabeela, Dongmei Li, Ashraf Ibrahim, Priya Uppuluri

## Abstract

A forward genetic screening approach identified orf19.2500, as a gene controlling *Candida albicans* biofilm dispersal and biofilm detachment. Three-dimensional (3-D) protein modeling and bioinformatics revealed that orf19.2500 is a conserved mitochondrial protein, structurally similar to, but functionally diverged from, the squalene/phytoene synthases family. The *C. albicans* orf19.2500 is distinguished by three evolutionarily acquired stretches of amino acid inserts, absent from all other eukaryotes except a small number of ascomycete fungi. Biochemical assays showed that orf19.2500 is required for the assembly and activity of the <u>N</u>A<u>D</u>H <u>u</u>biquinone oxidoreductase Complex I of the respiratory electron transport chain, and was thereby named *NDU1*. *NDU1* is essential for respiration and growth on alternative carbon sources, important for immune evasion, required for virulence in a mouse model of hematogenously disseminated candidiasis, and for potentiating resistance to antifungal drugs. Our study is the first report on a protein that sets the *Candida*-like fungi phylogenetically apart from all other eukaryotes, based solely on evolutionary “gain” of new amino acid inserts that are also the functional hub of the protein.

## Introduction

*C. albicans* biofilms are dynamic communities in which transitions between planktonic and sessile modes of growth occur interchangeably in response to different environmental cues. Biofilms growing on mucosal tissues or indwelling medical devices serve as localized reservoirs of highly drug resistant cells. Cells that disperse from this nidus into the systemic environment cause biofilm-associated disseminated infections (1, 2). Our previous reports have shown that biofilm dispersed cells are predominantly lateral yeast cells released from the hyphal layers of the biofilm (3). Phenotypically, biofilm-dispersed yeast cells have considerably better adherence to, and invasion of human tissues when compared to planktonic cells, and thereby are significantly more virulent than their free-living counterparts (3, 4). Global transcriptomic analysis of dispersed cells corroborated the virulence attributes, revealing expression of adhesins, invasins and secreted aspartyl protease genes, at levels similar to, or even statistically enhanced than parent biofilms (4). Interestingly, it was also found that the dispersed cells are transcriptionally reprogrammed before release, to acquire nutrients such as zinc and amino acids and to metabolize alternative carbon sources, while their biofilm-associated parent cells did not induce high-affinity transporters or gluconeogenetic genes, despite exposure to the same nutritional signals (4). Expression of genes required during starvation such as those encoding transporters, the TCA cycle and glyoxylate cycle components also implies that dispersed lateral yeast cells may have enhanced respiratory capacity over their metabolically dormant hyphal parents.

While regulatory networks governing *C. albicans* biofilm formation have been well-defined (5), hardly anything is known about the genes/proteins controlling biofilm dispersal. To date, *C. albicans* protein PES1 is the only molecular regulator that has been shown to control production of lateral yeast cells from hyphae, and to induce biofilm dispersal (4, 6). Thus, we embarked on a study to identify additional novel regulators of biofilm dispersal. Considering that dispersed cells are in a developmental phase distinct from the biofilm state, we hypothesized that some regulators may have a role in cellular metabolism or respiration.

Here, we report on the discovery of *NDU1*, a gene that encodes a mitochondrial protein required for the assembly and activity of the <u>N</u>A<u>D</u>H <u>u</u>biquinone oxidoreductase Complex I of the respiratory electron transport chain. Studies in *C. albicans* using gene deletion and complementation mutants revealed that *NDU1* is important for lateral yeast production and biofilm dispersal, and absence of *NDU1* triggers early biofilm detachment from its growth substrate. Our results further showed that *NDU1* is essential for respiration and growth on alternative carbon sources, potentiates resistance to antifungal drugs, is important for immune evasion, and full virulence in a mouse model of hematogenously disseminated candidiasis. Importantly, NDU1 protein has diverged significantly from other eukaryotic orthologues including the human orthologue NDUFAF6 (7); NDU1 protein harbors stretches of amino acid sequences acquired over evolution, that are uniquely specific only to Candida-like fungi, and can be the target for development of novel therapies.

## Results

### Loss of orf19.2500 abrogates *C. albicans* biofilm dispersal and induces early biofilm detachment

To identify potential regulators of biofilm dispersal, we performed forward genetic screening of several libraries of *C. albicans* mutants available through the fungal genetics stock center (FGSC) (Manhattan, KS). The libraries encompassed disruption mutants in *C. albicans* genes encoding transcription factors, kinases, cell wall integrity, and hundreds of other non-essential genes. Biofilms were developed from stationary phase cultures of each mutant on the MBEC Assay^®^ plates (Innovotech, Edmonton, Canada) which enables for high throughput screening of biofilm formation and dispersal (8). Of all the mutant strains that grew robust biofilms, two were isolated for their significant reduction in the frequency of biofilm dispersal. One strain was a mutant of the *C. albicans* major phosphodiesterase gene *PDE2*. The role of *PDE2* in cAMP-mediated control of lateral yeast production from hyphae has been previously published, and hence reduced dispersal from biofilms was expected (9). The other strain with abrogated biofilm dispersal was a mutant with deletions in an uncharacterized gene, orf19.2500.

Independent gene-deletion mutants of orf19.2500 (orf19.2500-/-) were constructed using a PCR-based gene disruption approach using small homology regions. In addition, complemented strains were constructed in which both alleles of orf19.2500 were reconstituted into the mutant (orf19.2500+/+). The ability of these mutant strains to develop biofilms were assessed in the 24-well polystyrene plates incubated under static conditions, or under flow of liquid medium on silicone elastomer (SE) material. Supernatant media from the static model, or media flowing over flow biofilms growing on SE were collected and quantified by measuring OD600 or by hemocytometer-based cell counts, respectively. Mutant biofilms displayed an overall 40%-50% decrease in biofilm dispersal compared to the wild-type (WT) or complemented strains in both static or the flow system (**Fig 1A**). Considering biofilm dispersal is a consequence of lateral yeast cells shed by biofilm hyphae in the surrounding media, top-most hyphal cells of flow biofilms were visualized under a microscope. Indeed, orf19.2500-/- hyphae showed a significant reduction in lateral yeast growth, compared to wild-type biofilm hyphae (**Fig 1B**), clarifying the reason for decrease in biofilm dispersal in the mutant.

**Figure 1.**
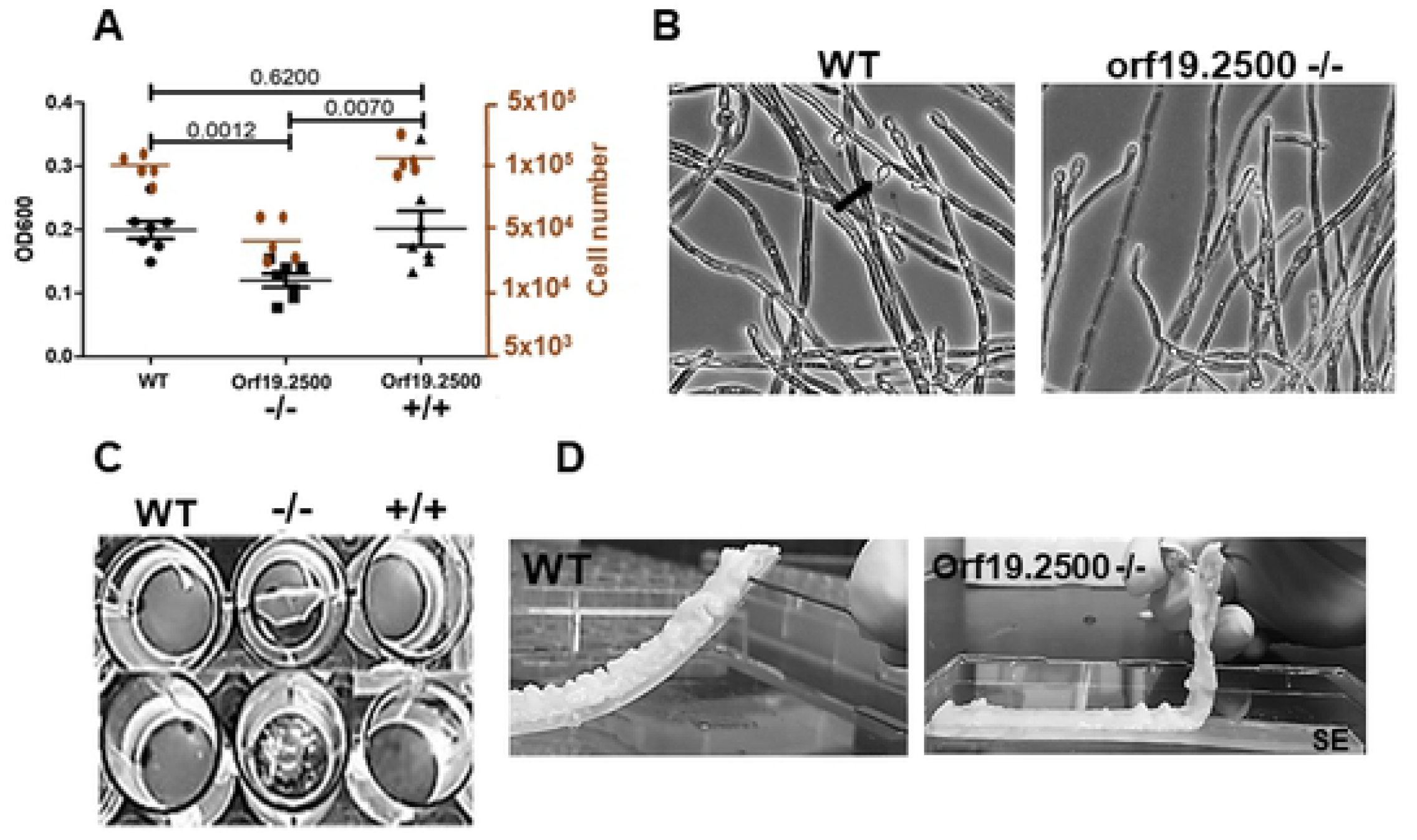
Extent of biofilm dispersal and attachment. (Note: Once its localization and function was determined, orf19.2500 was renamed as *NDU1* in later figures). *C. albicans* WT, orf19.2500 deletion mutant -/-, and revertant +/+ were allowed to develop biofilms using RPMI medium, under static and flow models for 24 h at 37°C. Cells released from the static biofilms growing on the pegs of the MBEC device were quantified using a spectrophotometer (OD600), and provided a measure of the extent of biofilm dispersal between the three conditions (black circles) (A). Values are average ±SEM; indicated p-values are measurements from seven independent replicates of the static biofilm model. Cells dispersed from biofilms grown in the flow model were collected after 24 h, counted using a hemocytometer, and plotted (orange ovals). Again, the p-value between WT and mutant or mutant and WT was <0.01 (not shown). Topmost layer of the biofilms were teased and imaged using a light microscope (40X mag), to visualize the extent of lateral yeast growth from mutant hyphae (arrow), versus the WT (B). Biofilms of the three strains were developed on 96 well microtiter plates under static conditions for 24 h, after which they were gently washed once to examine the robustness of their attachment to the well surface (C). Biofilms of the WT and orf19.2500 mutant were developed overnight on the surface of silicone elastomer under continuous flow of fresh RPMI at 37°C. The biofilms were gently teased away from the substrate to test the sturdiness of their attachment to the SE strips (D)

Under static biofilm induction, both orf19.2500-/- and +/+ developed biofilms comparable to wild-type, but only mutant biofilms detached early (16-20 h of growth) (**Fig 1C**). The layer of biofilm formed by the mutant strains either peeled off completely, or broke into pieces upon gentle washing of the biofilms with phosphate buffered saline. The detachment was not due to the inability of the mutant cells to adhere or form a robust biofilm, because orf19.2500-/- displayed comparable adherence to plastic and biofilm growth (data not shown). Similarly, under flow biofilm conditions, both wild-type and orf19.2500-/- developed robust biofilms; but while the wild-type biofilms were firmly attached to SE after 24 h of growth, mutant biofilms were easily displaced from the surface (**Fig 1D**).

### Orf19.2500-/- has a wild-type growth rate and morphology in glucose, but fails to grow on alternative carbon sources or in the presence of stressors

The fact that orf19.2500-/- was able to make robust biofilms indicated that it may not be deficient in growth or morphogenesis. We performed assays to measure the growth of the mutant in comparison to wild-type and complemented strains, under planktonic conditions. We found that in the first 24 h, the growth curves of orf19.2500-/- exactly overlapped with the other two strains, when grown in rich media containing 2% glucose (**Fig 2A**). In fact, the mutant and the wild-type controls displayed comparable growth rate and viability until day 4, after which the mutant cells gradually lost viability at significantly higher rates than the wild-type cells (**Fig S1A**). Visual examination of colonies grown on solid agar media, from 4 day-old cultures showed that the mutant cells were significantly smaller in size compared to wild-type cells, pointing to a defect in respiratory capacity post glucose exhaustion (**Fig S1B**). Microscopic visualization and measurement of hyphal lengths revealed no significant differences in hyphal growth and elongation between wild-type and mutant cells, and correspondingly no defect in their capacity to damage human vascular endothelial cells (**Fig S1C,D**).

**Figure 2.**
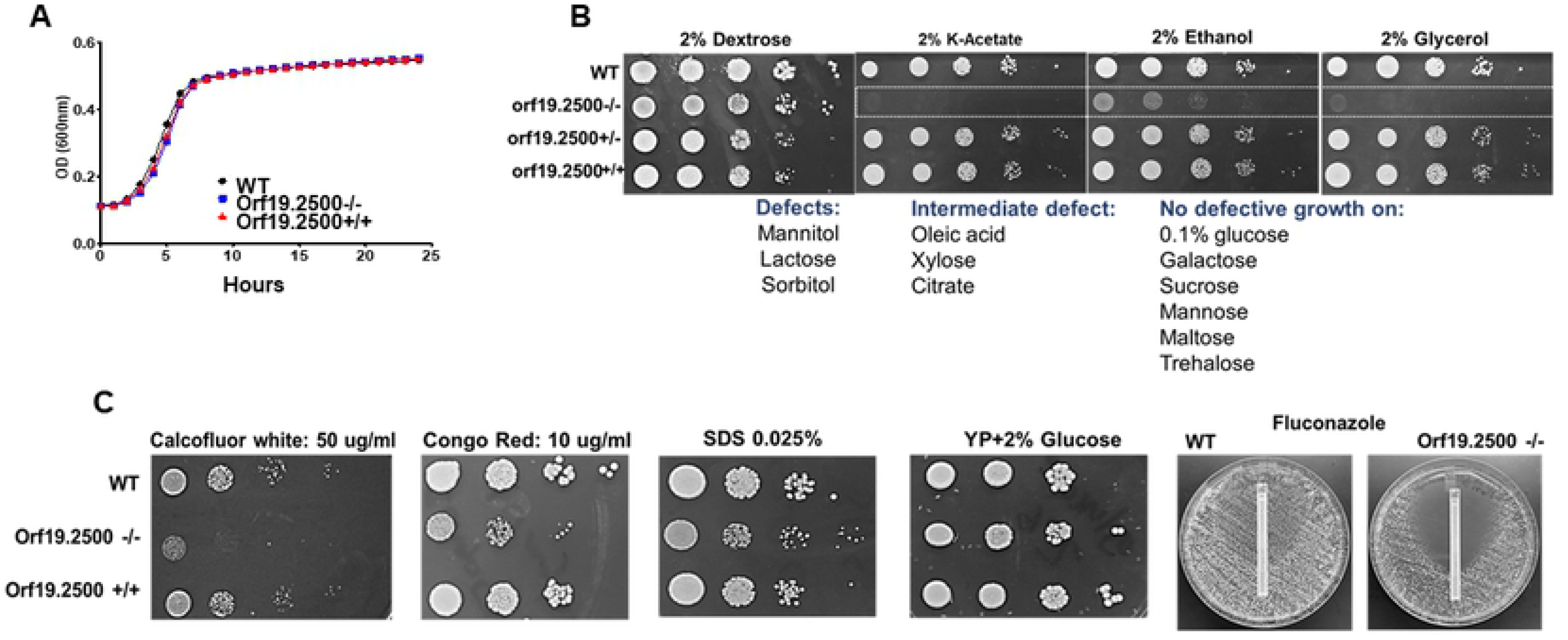
Pattern of growth on various carbon sources and stressors. The orf19.2500-/-, WT and revertant strain were grown in 10% YP+2% glucose media, and OD600 of the growth was measured temporally, using a spectrophotometer (A). Different concentrations (5 μl of 10^5^ cells/ml to 10^1^ cells/ml) of WT, orf19.2500 mutant -/-, heterozygote +/−, and revertant strain +/+ were spotted on solid YP media containing 2% of glucose, various alternative carbon sources or media containing cell wall/membrane stressors. Differences in extent of growth were visually noted (B,C).

It is well known that growth of petite sized colony of mutants is limited to fermentable carbon sources (10). As such, orf19.2500-/- grew robustly on media with glucose, but displayed severe growth defects on alternative carbon sources such as acetate, ethanol, glycerol (**Fig 2B**). The wild-type, heterozygote or the complemented strain did not exhibit this defect. We probed the extent of growth deficit of the mutant further, in the presence of cell wall and cell membrane stressors. Compared to the wild-type or the orf19.2500+/+ complemented strains, orf19.2500-/- disruption mutant was significantly more sensitive to growth on calcofluor white, congo red (cell wall stress), and SDS or fluconazole (cell membrane stress) (**Fig 2C**). In fact, on examination by staining with concanavalin A (stains cell surface mannans) or calcofluor white (stains cell wall chitin) followed by flow cytometry, the mutant strain displayed at least 40-50% reduced cell wall mannan or chitin content compared to the wild-type or complemented cells (**Fig S2A**). This deficiency was further corroborated by transmission electron microscopy, which displayed a strikingly thinner mannan layer, and at least 35% decrease in overall cell wall thickness in mutant cells versus the wild-type (**Fig S2B**).

To understand why the mutants were susceptible to cell membrane stressors, we investigated the membrane permeability of wild-type and mutant cells, as a measure of membrane integrity. Orf19.2500-/- mutant was significantly more permeable to fluorescein diacetate (FDA, a membrane intercalating dye), while wild-type cells completely blocked the membrane penetration of FDA, indicating that the cell membrane of the wild-type was healthier than that of the mutant (**Fig S2C**). As expected, the membrane disrupting antifungal drug fluconazole, which was used as a positive control, showed enhanced uptake of FDA in both wild-type and mutant cells. Since ergosterol is a major component of the *C. albicans* cell membrane (11), we questioned if ergosterol production was impaired in the mutant. Gene expression analysis of select *ERG* genes indicated that the orf19.2500-/- had a 2.7-fold increased expression of *ERG20* which is a putative farnesyl pyrophosphate synthase, required for both coenzyme Q and ergosterol biosynthesis. Most other *ERG* genes downstream of *ERG20*, solely important for ergosterol biosynthesis were downregulated >2-3 fold (**Fig S2D**), perhaps signifying the reason for higher membrane permeability and fluconazole susceptibility in the mutant.

### Orf19.2500 localizes to the mitochondria and plays a key role in functioning of Complex I of the mitochondrial electron transport chain (ETC)

In a quest to understand the function of this protein, we endeavored to unravel its cellular location. Attempts to tag one allele of orf19.2500 with a fluorescence tag were not productive due to a faint fluorescence signal, which although visible under the microscope, could not produce clear images for documentation. Because the expression levels of orf19.2500 were low, we constructed a tetracycline regulatable strain, which also harbored an mCherry marker right after the Tet-off promoter (orf19.2500/*ORF19.2500*-mCherry-tetO). In rich media containing glucose, there was an overexpression of orf19.2500-mCherry, and the protein in the cell fluoresced red. The red fluorescence completely overlapped with a stain that colors the mitochondrial matrix green, to provide an overall yellow colored mitochondrial localization (**Fig 3A**). Thus, the protein localized to the mitochondria in both yeast and hyphae.

**Figure 3.**
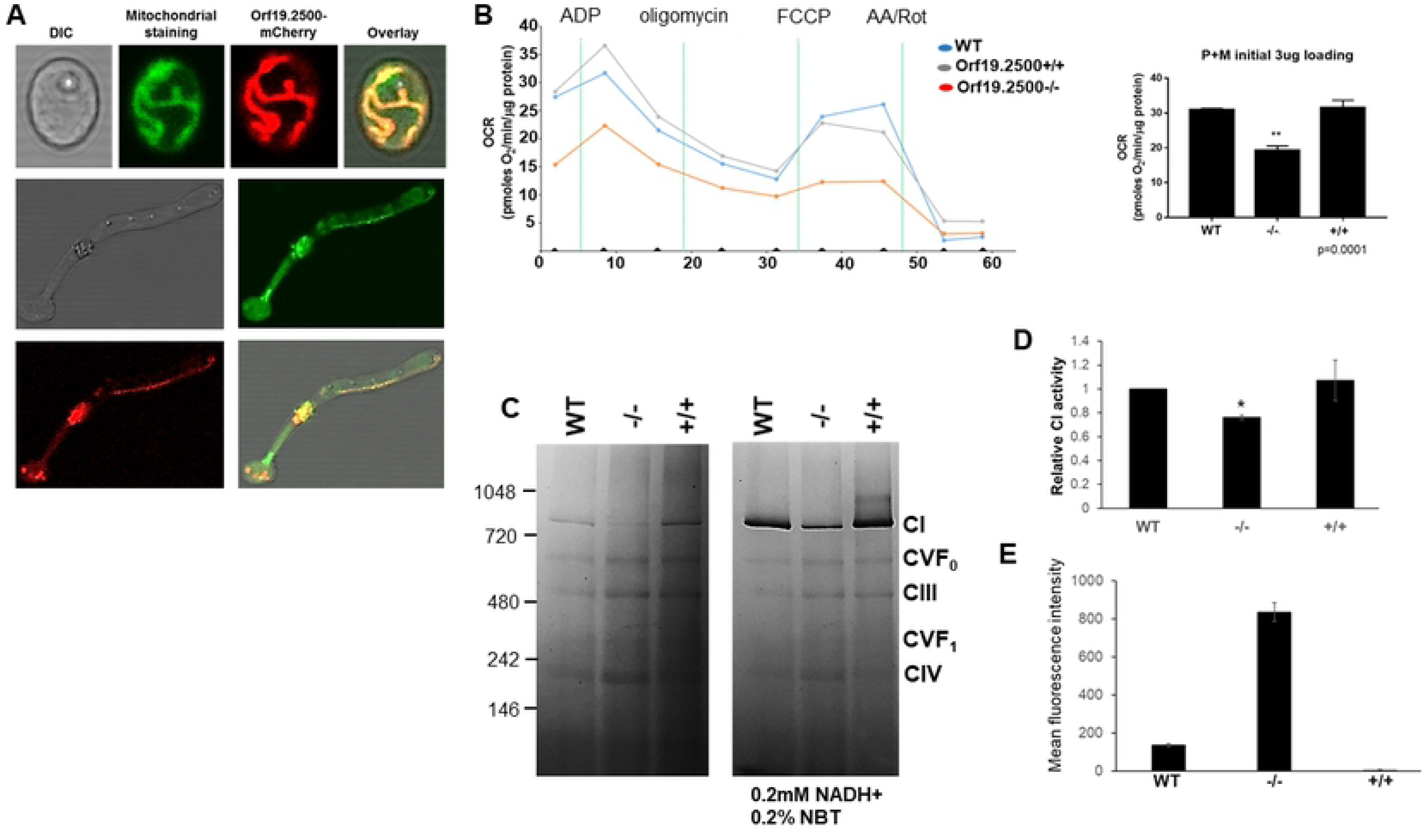
Organelle localization of orf19.2500 and measurement of its defect in respiration and mitochondrial complex stability. Orf19.2500 was engineered under a constitutively expressing tet-promoter, tagged with mCherry. Localization of the orf19.2500 was determined by staining both yeast and hyphal cells with a green fluorescent mitochondrial stain (Mitotracker green), to display the yellow overlap of colors in the mitochondria (A). The XF96 Analyzer was used to measure changes in mitochondrial bioenergetics by measuring the oxygen consumption rate (OCR) in freshly isolated mitochondria of WT, mutant, and revertant strains. One μg each of ADP, oligomycin, FCCP, and antimycin A and rotenone were added at the indicated points. The maximal respiratory capacity was quantified (B). Values are means±SEM. **(p<0.01 mutant vs. WT and revertant; measurements from six independent isolations). BN-PAGE electrophoresis of equal quantities of total mitochondrial proteins of *C. albicans* WT, orf19.2500-/- and orf19.2500+/+ cells, stained with Coomassie to reveal respiratory complexes CI to CIV (C). The molecular markers indicated to the left are NativeMark (unstained protein standard; Invitrogen). The in-gel enzyme activity of CI in BN-PAGE was assayed within 60 min after incubating the gel in reaction medium (0.1 M Tris-HCL, pH 7.4, 0.2 mM NADH as a substrate, and 0.2% NBT). Specific activity of CI in mitochondria from mutant and complemented strains were quantified and plotted relative to the CI activity in wild-type mitochondria (* = p<0.05) (D). ROS activity in wild-type, mutant and complemented strains were measured by staining the cells with MitoSox red, and measuring fluorescence intensity was measured by flow cytometry and plotted (E).

The fact that orf19.2500 mutant could not grow on alternative carbon sources indicated a respiratory defect likely due to a faulty electron transport chain (12). To test this hypothesis, we carried out Seahorse assays to test the respiratory prowess of the isolated mitochondria of the mutant strain, in the presence of Complex I (CI) substrates (pyruvate + malate) (**Fig 3B**). Compared to the mitochondria from the wild-type or the complemented strain, the orf19.2500-/- mutant mitochondria showed an overall significant decrease in respiratory capacity, measured by a 30% decrease in oxygen consumption rate. This was not the case when a Complex II (CII) substrate (succinate in presence of rotenone) was used; Mitochondria from all three strains displayed equivalently robust CII activity (**Fig S3A**).

To further test if CI was impacted in the mutant strain, we performed a Blue native PAGE (BN-PAGE) analysis, in which the five different complexes of the ETC from isolated mitochondria (of each strain respectively), were separated by electrophoresis, and CI activity tested. We determined that CI is reduced by 40 to 50% in orf19.2500-/- compared to the gel density ratios of CI/CIII and CI/CV to wild-type or orf19.2500+/+ cells by ImageJ (**Fig. 3C**). In addition, the *in situ* assay of CI NADH dehydrogenase enzyme activity demonstrated that mutant strain correspondingly had reduced enzyme activity than the wild-type or reconstituted mitochondria (**Fig. 3C**). Quantitative measurement of enzymatic activity independently confirmed a ~30% decrease in CI in the mutant compared to the wild-type strains (p<0.05) (**Fig 3D**). Thus, the seahorse assays (**Fig. 3B**), reduced assembled CI and its enzymatic activity (**Fig 3C,D**), support the hypothesis that orf19.2500 is important for CI activity in *C. albicans*. Based on its role in mitochondrial oxidative phosphorylation and its NADH dehydrogenase activity, Orf19.2500 was renamed as *NDU1* for NADH dehydrogenase of Complex I.

Since CI is the major donor to the proton gradient (13), we posited that a reduction in its activity would affect the mitochondrial membrane potential (μψM) Wild-type, mutant and complemented strains were grown overnight in YP+2% glucose, and then sub-cultured in media containing either glucose or acetate as a carbon source. After 2 h of growth, cells were treated with JC1, a dye used as an indicator of mitochondrial membrane potential. Compared to wild-type and complemented strains, *NDU1* mutant cells had significantly higher mitochondrial membrane depolarization, as measured by the intercalation of JC1 dye, and the shift in the green FITC fluorescence. However, this reduction in membrane potential was found only under nutritional stress, such as in the presence of the alternative carbon sources of acetate (**Fig S3B**) or sorbitol (data not shown), and not during growth on glucose.

CI is responsible for most cellular ROS production in mitochondria (13), and impairment in CI activity often results in oxidative stress. The accumulation of mitochondrial ROS was determined by measuring the superoxide levels with MitoSOX Red dye, which is specifically targeted to mitochondria in live cells. Oxidation of MitoSOX Red reagent by superoxide produces red fluorescence, which is quantified by flow cytometric analysis and correlated with the amount of ROS present in the mitochondria. *NDU1* deletion led to an elevation in the mitochondrial superoxide levels, which upon quantification of flow cytometric data, revealed a greater than 6-fold increase in MitoSOX staining in *NDU1* mutant cells versus the wild-type or complemented strains (**Fig 3E)**.

### *NDU1* is hyper-susceptible to neutrophil killing and avirulent in a mouse model of hematogenously disseminated candidiasis

Considering that *NDU1* mutants are unable to grow on alternative carbon sources, we hypothesized that they would have difficulty surviving in the nutritionally deprived environment of innate immune cells (14). To test this, we determined the killing ability of these mutant strains by human neutrophils. Within 45 min, neutrophils had engulfed yeast cells of all three strains. By 90-150 min, *C. albicans* wild-type as well as the complemented strain developed germ tubes, and were able to destroy the immune cells (**Fig 4A**). In contrast, the *NDU1* null mutant remained as engulfed yeast cells inside the neutrophils, and by 3 h were eventually killed in significantly (2-fold) higher numbers than wild-type or *NDU1* complemented strains (**Fig 4B**).

**Figure 4.**
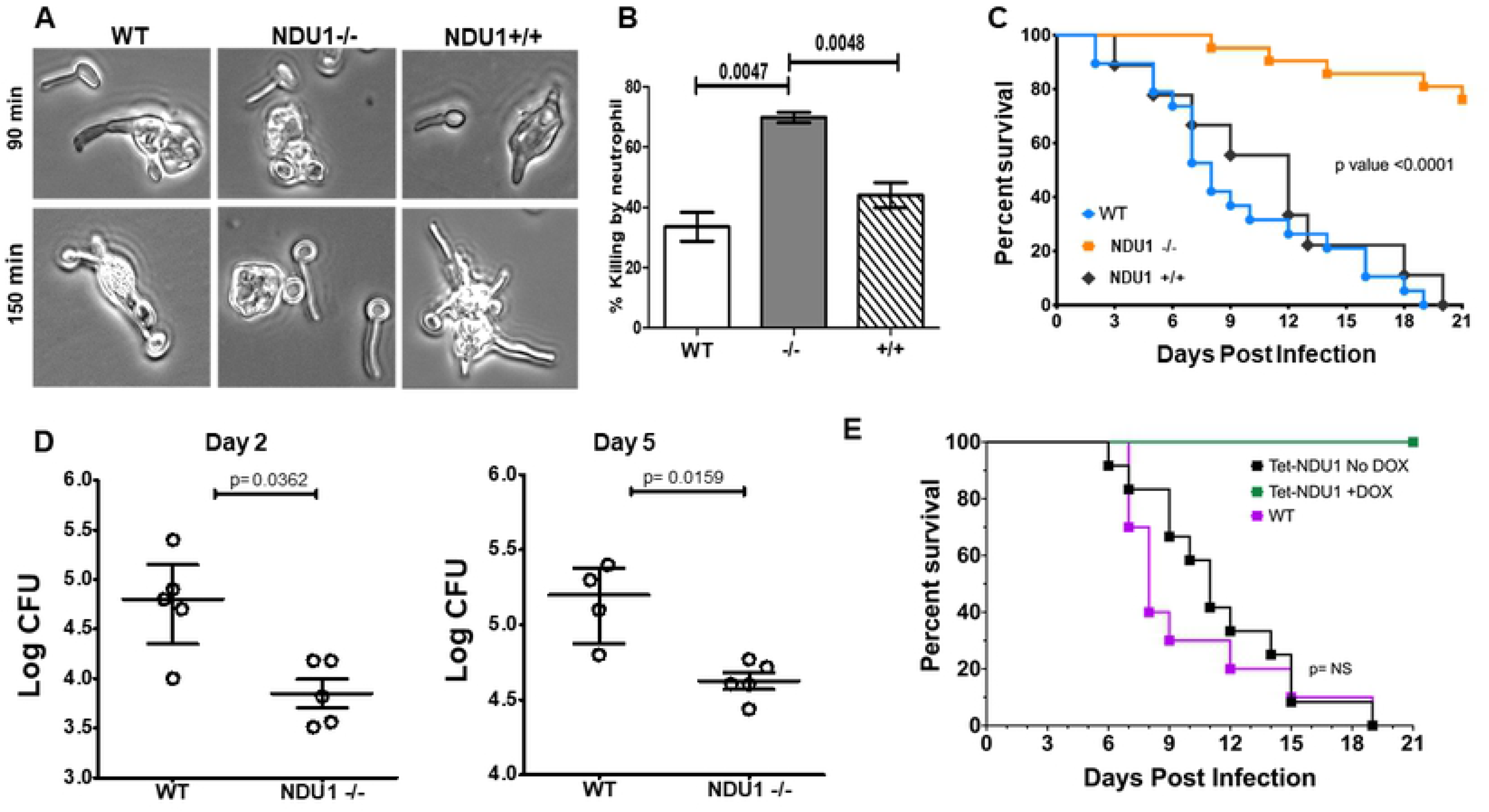
*Note: orf19.2500 is referred as NDU1 from this figure on*. Susceptibility to neutrophils and defective virulence of NDU1 mutant. Yeast *C. albicans* WT, NDU1-/- and NDU1+/+ cells grown overnight were incubated along with primary human neutrophils (3:1 MOI), for 3 hours, and the extent of phagocytosis was visualized by confocal microscopy at 90 min and 150 min (A). The extent of yeast cell killing by neutrophils was quantified by CFU measurement at 3 h (A). Data are mean + SEM from three biological replicates. CD1 outbred mice were infected via tail vein with 2.5×10^5^ cells of WT, NDU1-/- and NDU1+/+ (10 mice each × 2 replicates) and the impact of disseminated candidiasis on the overall survival of mice was monitored for 21 days. *P < 0.0001 mutant vs. wild-type and revertant, log-rank test (B). Mice were sacrificed on day +2 and +5 relative to infection, and their kidneys processed for tissue fungal burden, determined by plating on solid media. Data are median ± interquartile range. *P < 0.05 mutant versus wild-type, Wilcoxon rank-sum test (C). Survival curves for the different groups of mice (10 animals per group) infected with the *C. albicans* WT isogenic parental strain or with the *C. albicans* tetO-*NDU1*/*ndu1* strain in the presence or absence of doxycycline (DOX). Statistically significant differences were measured between the comparison groups of mice (D).

The inability of *NDU1*-/- to evade the immune system, translated predictably into avirulence in a hematogenously disseminated mouse model. While 100% of the mice succumbed to infection by the wild-type or complemented strains within 21 days, >80% of the mice infected with *NDU1*-/- null mutant survived the infection (**Fig 4C**). This striking defect in survival was corroborated with ~ 0.5-1.0 log reduction in kidney fungal burden of mice infected with the *NDU1-/- vs.* those harvested from mice infected with the wild-type and collected 2 or 5 days post-infection (**Fig 4D**).

We also studied the virulence of the generated mutant strain at 10-fold higher infectious dose of 2.5×10^6^ cells. Interestingly, while mice infected with wild-type and complemented strains succumbed early to the infection within 7 days, ~80% of mice infected with the *NDU1-/-* mutant survived the infection (**Fig S3C**), thereby mimicking the survival of mice infected with the lower infectious dose (**Fig 4C**).

The fact that the mutant strain did not cause virulence could likely be attributed to their early susceptibility to phagocytes, or their defective long-term sustenance in a glucose-impoverished environment *in vivo*. To test this further, we constructed a *C. albicans* strain, wherein one allele of *NDU1* was deleted while the other was placed under a tetracycline-regulatable promotor (*NDU1Δ*/*NDU1*-tetO). The expression of *NDU1* could be increased or decreased based on the presence or absence of doxycycline (DOX) in the growth milieu. For virulence studies, one set of mice were infected via tail vein with WT, and two other sets with the *NDU1*-tetO strain. Mice were fed with plain water for 24 h after infection, to enable unrestrained dissemination, after which DOX was added (at 24 h after infection) to the drinking water of one set of *NDU1*-tetO infected mice (to deplete expression of *NDU1*) and to the set infected with WT (DOX control). The third set of mice were fed continuously with water without DOX (for overexpression of *NDU1*). As clearly seen in **Fig 4E**, sustained depletion of *NDU1 in vivo* due to DOX in systemic circulation translated into 100% mouse survival rate, while the WT and overexpression strains demonstrated similar levels of lethality in mice.

### *NDU1* 3D structure has characteristics of a dehydrosqualene synthase, and is homologous to human NDUFAF6

We predicted that the key to identifying NDU1 protein function lay in unraveling its 3D structure. The NDU1 sequence was submitted to MitoProt II for analysis (15); a mitochondrial targeting sequence of 15 amino acids was predicted to be removed with high probability (0.9124), suggesting the mature NDU1 (mNDU1) protein begins at Asn16. Structural models for NDU1 were generated by submitting the mNDU1 protein sequence to Phyre2 (V 2.0), a protein homology recognition engine that uses profile-profile matching algorithms (16). A model for NDU1 named c5iysA was generated with 100% confidence by threading the NDU1 sequence onto chain A of 5IYS. 5IYS is the crystal structure of the *Enterococcus hirae* dehydrosqualene synthase in complex with two molecules of the substrate analog, farnesyl thiopyrophosphate (FPS) (**Fig 5A**). An overlay of c5iysA and 5iys has an RMSD of 0.270 Å between 253 pruned atom pairs; an excellent match, especially over the core regions. The two molecules of FPS are bound in a large “pocket” (2087 Å^3^) in 5iys (**Fig S4A**). The c5iysA model (NDU1 threaded onto 5iys) also has a large pocket as determined by Castp (17) (2332 Å^3^) (**Fig S4B**). The pocket is larger than that in 5IYS but has a somewhat different shape and cannot accommodate the FPS molecules exactly as positioned in 5IYS. Nonetheless, the FPS lipid chains are highly flexible and can likely adapt to the c5iysA pocket. The second model predicted by Phyre2 was c4hd1A, which is modeled on 4hd1 which is a squalene synthase from *Alicyclobacillus acidocaldarius* (**Fig S4C**). Overlay of c4hd1A (green) and 4hd1 (purple) yielded an excellent RMSD between 245 pruned atom pairs as 0.262 angstroms. Thus, NDU1 is a predicted squalene/phytoene synthase (pfam: PF00494).

**Figure 5.**
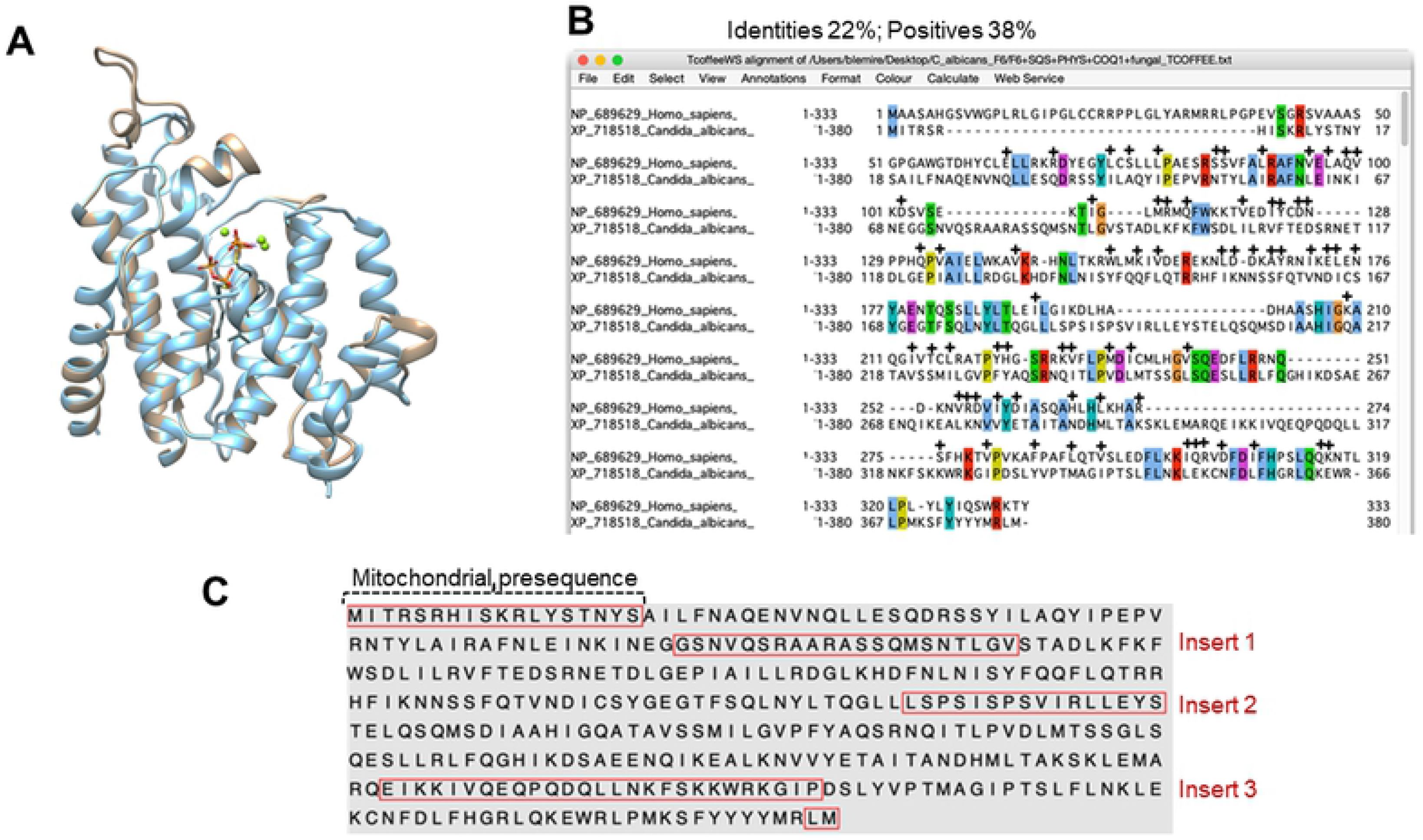
Overlay of template and NDU1 models. **The** Phyre2 model of NDU1, c5iysA (blue) is overlaid onto the structure 5iys (tan) with an RMSD of 0.270 Å between 253 atom pairs. Two molecules of the substrate analog, farnesyl thiopyrophosphate (FPS) and three Mg^++^ ions (green) are bound in the 5iys active site cavity. (A). BLAST alignment of *C. albicans* NDU1 with human NDUFAF6. Colored amino acids represent identity, while + indicates positives (B). Protein sequence of NDU1 with the highlighted mitochondrial presequence, and three inserts acquired over evolution, which sets *C. albicans* NDU1 apart from its human orthologue (C).

We also unraveled that *C. albicans* NDU1 has a human orthologue NDUFAF6, the NADH:ubiquinone oxidoreductase complex assembly factor 6, which shares approximately 22% identity (38% similarity) (**Fig 5B**). Location of specific NDU1 residues (red) with identities to NDUFAF6 (grey) are displayed in a structural schematic in (**Fig S4D**). NDUFAF6 is considered a member of the Isoprenoid_Biosyn_CI superfamily, which generates tens of thousands of isoprenoid metabolites, including sterols, heme, dolichol, carotenoids and ubiquinones.

### *C. albicans* NDU1 has evolutionarily acquired amino acid inserts unique to CTG clade fungi

Protein sequence alignments and 3D homology modeling between NDU1 and NDUFAF6 further revealed that NDU1 is a longer protein (380 *vs.* 333 amino acids of NDUFAF6), and it has extra stretches of amino acid inserts, that are depicted as gaps in the human NDUFAF6 sequence (**Fig 5B**). Specifically, NDU1 has four prominent amino acid inserts that are evolutionarily acquired within its protein sequence (**Fig 5C**). The part of sequence highlighted at the N-terminus is the predicted mitochondrial targeting sequence, which is truncated upon import to the mitochondria, hence is irrelevant to function. The red-boxed inserts 1, 2 and 3 are unique to NDU1. These inserts are not modeled in the c5iysA model from PHYRE2, as they are not present in the 5IYS template. The mNDU1 sequence was submitted for modeling to iTASSER, which employs threading template identification and iterative modeling to model the entire sequence (18). On the iTASSER model, the three inserts lie in loops on the surface of the protein model (**Fig 6A**). In fact, when visualized on a surface model, insert 1 is located at the opening of the NDU1 pocket. The pocket/cavity is formed by long alpha helices packing together, and inserts 2 and 3 were modeled to the bottom of the V-shaped pocket cavity, between those alpha helices (**Fig 6B**). Additionally, the latter two inserts were found close together in 3D space, enough to be in contact with each other.

**Figure 6.**
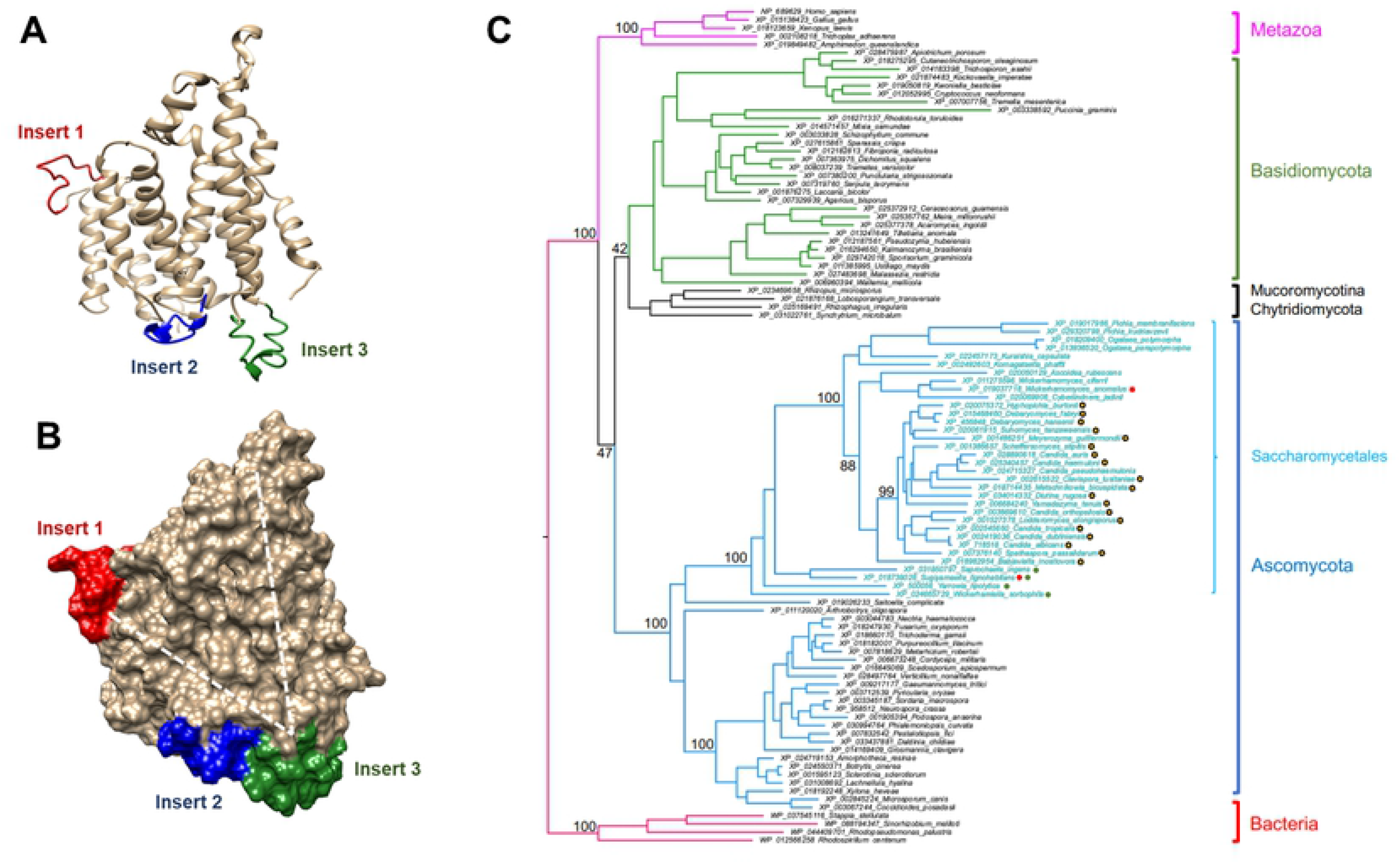
Location of the three insertion sequences of NDU1 and their phylogenetic uniqueness. The unique insertion sequences of NDU1 were localized on ribbon and on surface representations of the iTASSER model (A). Note how insert 2 (blue) and 3 (green) lie to the bottom of the V-shaped cavity highlighted by dotted lines (B). A maximum likelihood tree of selected eukaryotic and prokaryotic NDU1 orthologs emphasizes the highly restricted distribution of the three insert sequences. This tree has 4 bacterial (red), 5 metazoan (magenta) and 93 fungal orthologs, with branches colored green for Basidiomycota, black for Chytridiomycota and Mucoromycotina, and blue for Ascomycota. Tips belonging to the order *Saccharomycetales* are colored cyan. Bootstrap support values for selected nodes are given. Proteins belonging to the CTG clade of yeasts are noted with a yellow star. All *Saccharomycetales* proteins have insert 2; two are completely missing insert 1 (red circles); 4 are completely missing insert 3 (green circles).

To investigate the phylogenetic distribution of the insertion sequences in orthologous proteins, we performed a Psiblast search using *C. albicans* NDU1 (XP_718518) as query against the RefSeq (release 200; 2020/05/04) database. Putative bacterial and eukaryotic orthologs of NDU1 with evalues < 1e-30 were aligned with MAFFT (19), a multiple sequence alignment software, and a phylogenetic tree was generated using IQ-TREE (20). The tree was used to manually reduce the number of sequences outside the *Saccharomycetales*, while retaining phylogenetic diversity. The final tree contains 102 putative NDU1 orthologs and uses the bacterial orthologs as an outgroup: 4 bacterial (red), 5 metazoan (magenta) and 93 fungal orthologs, with branches colored green for Basidiomycota, black for Chytridiomycota and Mucoromycotina, and blue for Ascomycota. Only the Saccharomycetales (in particular the CTG-clade yeasts noted with a yellow star) were found to contain all three inserts. This group had longer branches and were well separated from the other groups, indicating greater divergence in the NDU1 sequences over evolution. Interestingly *Saccharomyces cerevisiae* and *Candida glabrata* are not included in the tree because they do not have CI, and hence lack *NDU1* orthologues (**Fig 6C**).

To elaborate further, based on the presence or absence of the three insertion sequences, the phylogenetic tree was clearly divided into three groups (**Fig S4E**): group 1 colored blue contained Saccharomycetales and *Candida* like CTG-clade fungi (CTG-clade), group 2 in black which had other fungi, and group 3 colored pink represented sequences from bacteria and eukaryotes. Group 1 had longer branches and were well separated from group 2 and 3 sequences. Also, only group 1 had all three insert sequences. In contrast, group 2 had no insert 1 or insert 2 and had a different insert 3, while group 3 were lacking in all the inserts (**Fig S4E**). Overall, our analyses showed that the *C. albicans* mitochondrial protein NDU1 has structures distinct to CTG-clade proteins, and these inserts may be functionally important for enzymatic activity or protein-protein interactions, distinct for *Candida* spp.

### Expression of the human NDUFAF6 does not complement *C. albicans* NDU1 defect, while insert 2 and 3 are the functional hub of NDU1

Since NDU1 was important for immune evasion, drug resistance and virulence in *C. albicans*, we questioned if it would represent a viable target for antifungal drug development. Specifically, we probed the functional diversity of NDU1 compared to its human counterpart NDUFAF6. Hence, we first inspected if heterologous expression of human NDUFAF6 in *C. albicans* could provide a gain of function in the NDU1 mutant strain. The entire codon optimized ORF of NDUFAF6 was expressed constitutively under a tet-regulatable promoter. Importantly, the expressed protein also harbored a GFP-tag. NDUFAF6-GFP localized correctly to the mitochondria of both yeast and hyphae (**Fig 7A**), and Western blotting of *C. albicans* total protein with anti-GFP antibodies yielded the correct size NDUFAF6 protein (~63 kDa; NDUFAF6 38kDa+GFP 25 kDa) (**Fig 7B**). We tested if expression of the human orthologue could revert *C. albicans* NDU1 mutant growth defect on acetate. As expected, the control *NDU-/-* mutant did not grow on acetate, glycerol, sorbitol or ethanol, while strains overexpressing the full length *C. albicans NDU1* (tet-regulatable NDU1; OE) grew robustly on the non-fermentable carbon containing media (**Fig 7C**). We also found that human NDUFAF6 expression did not restore growth of the mutant on 2% potassium acetate, or other alternative carbon sources such as ethanol, glycerol or sorbitol (**Fig 7C**), indicating that the human protein could not revert the functional defect of NDU1 on non-fermentable carbon sources.

**Figure 7.**
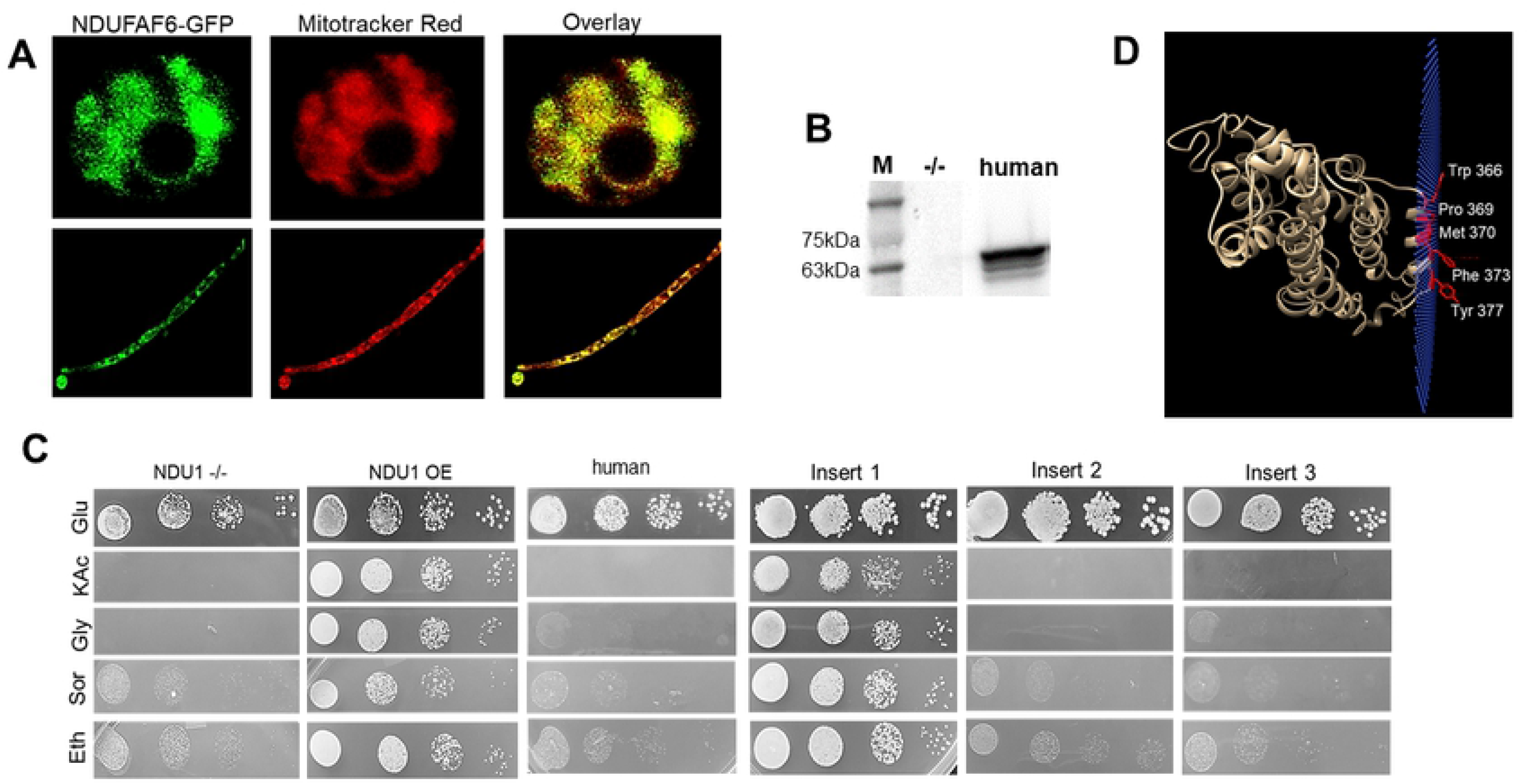
Expression and localization of human NDUFAF6 in *C. albicans.* Entire ORF of GFP-tagged human NDUFAF6 was expressed in *C. albicans* NDU1 mutant, and found to be localized to mitochondria, as visualized by GFP overlapping with a mitochondrial stain, in both yeast and hyphae (A). Detection of the GFP-tagged human NDUFAF6 protein by Western blotting using an anti-GFP antibody. M=marker, second lane is total protein from NDU1 mutant (negative control), lane 3 is the detected 63 kDa human NDUFAF6-GFP fusion protein (B). C. albicans NDU1 mutant expressing NDUFAF6 grows on glucose but not on 2% acetate (B). Individually expressed entire ORF of NDU1 (NDU1 overexpression strain), or NDU1 lacking each insert, in the NDU1 mutant, were screened for their ability to grow on alternative carbon sources. The NDU1 mutant was used as the parent control strain for the experiment (C). OPM server predicts NDU1 is a peripheral membrane protein (plane of blue spheres) and identifies five membrane-embedded amino acids (D).

To further validate this, we probed if the three insert sequences unique to *C. albicans* NDU1 had a role to play in mitochondrial function determined by growth on non-fermentable carbon sources. As done with the human protein, we expressed the entire ORF of *C. albicans* NDU1 minus the individual inserts, in the *C. albicans NDU1* mutant. Similar to the *C. albicans* mutant expressing the human protein, mutants expressing the mutated *C. albicans* NDU1 constructs expressed proteins that localized to the mitochondria, and were expressed at the right size, as evaluated by fluorescence microscopy and Western blots, respectively (data not shown). Interestingly, we found that strains expressing NDU1 with a truncated insert 1 grew as good as the OE strain, indicating that this area was dispensable for protein function. In contrast, mutants with individually truncated insert 2 or 3, maintained the growth defect on alternative carbon sources, highlighting their importance to the function of NDU1 (**Fig 7C**). All strains grew robustly on glucose containing media. As a control, we also randomly truncated two stretches of NDU1 (corresponding to ~30 amino acids each) in locations other than the inserts. This construct when expressed in *C. albicans* behaved just like insert 1 (data not shown), providing further proof that insert 2 and 3 are likely the prime areas for enzymatic activity, or sites of interaction of NDU1 with other proteins in the mitochondria.

### NDU1 is a peripheral membrane protein

Isoprenoid substrates and products are hydrophobic, and predicted to be localized to the membrane. NDU1 was not found to have predicted transmembrane segments (using transmembrane prediction servers such as TMHMM Server v. 2.0 or PRED-TMR of the SwissProt database). We then submitted the iTASSER model to the OPM server (Orientations of Proteins in Membranes), which determines the spatial position of membrane boundaries for integral and peripheral membrane proteins (21). OPM optimizes the free energy of protein transfer (ΔG_*transf*_) from water to a membrane environment. For peripheral membrane proteins ΔG_*transf*_ varies between −15 and −1.5 kcal mol^−1^. OPM strongly suggested that NDU1 is a peripheral membrane protein with optimal energy of −7.6 kcal/mol, and 5 residues of the C-terminal helix (Trp356, Pro369, Met370, Phe373 and Tyr377) were predicted to be membrane embedded (**Fig 7D).**The model also predicted the NDU1 cavity to lie above the mitochondrial membrane (**Fig S4F**).

## Discussion

Biofilm formation and dispersal are determinants of virulence in *C. albicans*. We used genetic, bioinformatic and biochemical approaches to identify *NDU1*, a gene that encodes a mitochondrial protein that is evolutionarily divergent from other eukaryotic orthologues. *NDU1* affects biofilm dispersal and detachment, helps resist phagocyte killing, and is important for full virulence of *C. albicans* in mice. The most compelling phenotype of *NDU1* null mutant is its inability to grow on non-fermentable carbon sources or in advanced stationary growth when glucose is depleted. In *S. cerevisiae* and *C. albicans*, respiration-deficient mitochondrial mutants are unable to grow on non-fermentable carbon sources (22), and form petite colonies because their cell division rates are lower than that of the normal cells (23). Consistent with these observations, and as predicted by the N-terminus mitochondrial targeting sequence (**Fig 5C**), the GFP-tagged *NDU1* localized to the *C. albicans* mitochondria, signifying that it may have functions in cellular respiration.

Loss of *NDU1* additionally lead to a striking reduction in *C. albicans* cell wall thickness due to prominent reduction in chitin and cell surface mannan content rendering them hypersusceptible to cell wall perturbing agents. This reduction in cell wall morphology was appreciated only after 48 h to 3 days of growth, around the time glucose was exhausted from the growth medium. As has been previously demonstrated by global gene expression profiling and biochemical studies, *C. albicans* cell wall and mannoprotein synthesis requires energy, most of which is provided through oxidative phosphorylation by the mitochondrial electron transport chain (24–27). Perhaps then, it was the defective growth phenotype of this mutant on alternative sources leading to cell wall alterations, which contributed towards the sub-optimal biofilm phenotype. Inner layers of a mature biofilm are nutrient starved and hypoxic (28), and respiration is important for survival of these cells (29). Premature detachment of biofilms formed by the mutant cells could prospectively be the consequence of loss of viability at the bottommost nutritionally disadvantaged layers of the biofilm, coupled with an early loss of adhesion to substrate due to alterations in the cell wall architecture. Analysis of expression levels of genes and proteins involved in viability or adhesion in the innermost cells of the mutant biofilms versus the wild-type biofilms will help understand the role of *NDU1* in maintaining the biofilm. Likewise, we have recently reported that when compared to planktonic or biofilm cells, biofilm-dispersed lateral yeast cells are metabolically rewired, expressing at significantly high levels, genes important for respiration, and nutrient assimilation (4). Thus, loss of *NDU1* leading to reduced biofilm dispersal could be because of deficient respiratory activity, due to which energy required for production of new daughter cells from quiescent biofilm hyphae is lacking.

Energy in the cell is generated in the form of ATP, and Crabtree negative organisms such as *C. albicans* rely upon oxidation of substrates via the mitochondrial tricarboxylic acid (TCA) cycle to generate ATP even in the presence of glucose (30, 31). In the classical respiratory pathway, complexes of the electron transport chain (CI, CIII, and CIV) pump H^+^ ions to the intermembrane space. This creates a membrane potential (ΔψM) which is used by the ATP synthase to synthesize ATP (32). Several lines of evidence including using sea horse assays with CI substrates, *in vitro* activity and *in situ* gel analysis, measurement of mitochondrial membrane potential, and measurements of ROS were used to show that the major dysfunction of *NDU1* mutant was due to a defect in CI. A reduction in CI activity is harmful to the cells because it is the hub of all energy provided by oxidative phosphorylation, and is the major ROS-generating unit in the mitochondria (13). Dysfunction of CI destabilizes membrane potential and triggers ROS release, which is often fatal to the cell. *NDU1* null mutant was defective in both maintaining an intact membrane potential, and keeping ROS under check.

*NDU1’*s shortcoming in utilizing alternative carbon sources and its disrupted cellular architecture caused mutant cells to be significantly more susceptible to primary human neutrophil cells. Indeed, innate immune cells are nutrient starved, and only those cells capable of growing under respiratory stress can combat these host cells (14). In fact, *NDU1* mutant was strikingly avirulent in mice, with kidney organ burden of at least a log lower than mice infected with the wildtype strains. This defect in virulence *in vivo* was not due to a general growth defect of the mutant because between day 2 and day 5 of infection, the number of infected cells in the target organ, kidneys, increased by 5-fold in both mutant as well as the wild-type strain (**Fig. 4D**). This virulence defect in the mutant was confirmed when *NDU1* expression levels were controlled *in vivo* (**Fig. 4E**). It is well known that the integrity and function of mitochondria are essential to the virulence of *C. albicans*. Mutations affecting any one of a number of mitochondrial functions, including mitochondrial ribosome synthesis, mitochondrial transcription or genome maintenance, protein import, or functioning of the electron transport chain (ETC), result in *C. albicans* virulence defect (12, 26, 27, 33, 34).

Our results on NDU1 and its role in the CI-related activity is reminiscent of another well-characterized CI protein in *C. albicans*, GOA1. *GOA1* deletion mutants fail to make complex I, resulting in reduced respiration, and multiple deficits on alternative carbon sources, cell wall alterations, enhanced sensitivity to killing by neutrophils, and reduced virulence in a murine model of disseminated disease (25, 32, 35, 36). However, the greatest difference between *NDU1* and *GOA1* is in their differential ability to undergo morphogenesis and form a biofilm. *GOA1* mutants cannot make hyphae or develop a biofilm (35), while *NDU1* is adept at both fronts. This indicates that *NDU1* might have regulatory functions different from *GOA1*. Interestingly, orthologues of *GOA1* are restricted to members of the “CTG clade” of fungi, whose members decode CTG codons as serine rather than leucine (35, 37), suggesting that it represents a lineage-specific mitochondrial adaptation. Given that significant differences exist in CI among various lineages, and its important role in the pathobiology of *C. albicans*, an in-depth understanding of CI is warranted.

The *C. albicans* CI is believed to consist of 39 proteins, and while several of these proteins are conserved in other eukaryotes, a few such as GOA1 are unique to the CTG clade of which *Candida* species is a part (12, 36). NDU1 however has orthologues in other eukaryotes including humans. All fungal orthologues of NDU1 can be grouped into one clade and are monophyletic, meaning they inherited the gene from the same ancestor. The predicted structure of NDU1 is completely helical with a large central cavity that can accommodate 2 farnesyl pyrophosphate molecules. Three dimensional modeling combined with bioinformatic and phylogenetic analysis revealed that NDU1 belongs to the family of dehydrosqualene synthases. Within this superfamily NDU1 belongs to the family of Trans-Isoprenyl Diphosphate (Pyrophosphate) Synthases (Trans_IPPS; CDD: cd00867) and specifically the head-to-head family (Trans_IPPS_HH; CDD: cd00683) of synthases, that catalyze the condensation of farnesyl or geranylgeranyl diphosphates to form squalene of cholesterol biosynthesis, or phytoene of carotenoid biosynthesis (38). When a phylogenetic tree (adapted from (7)) with 50 Trans_IPPS domains was generated (**Fig S5A**), three monophyletic groups emerged: Cluster I proteins are putative NDU1 orthologues in fungi (green) and other eukaryotes and prokaryotes (pink); cluster 2 proteins that are phytoene synthases (PHYS) and squalene synthases (SQS) and cluster 3 which are the prenyl diphosphate synthases called COQ1 (coenzyme Q). The eukaryotic cluster 1 proteins from humans and drosophila have been experimentally localized to the mitochondrial inner membrane (39). Likewise, NDU1 is predicted to be a peripheral membrane protein, embedded into the membrane via five amino acid residues of the C-terminus.

NDU1 is 22% identical to its human orthologue NDUFAF6, with ~38% overall similarity between the two proteins. The Trans_IPPS_HH proteins typically have two DxxxD motifs involved in substrate binding and the coordination of catalytically important Mg++ ions (40). NDU1, NDUFAF6 and their orthologues do not have these motifs, suggesting the possibility that they have lost the ability to function as squalene/phytoene synthases (7). Besides, humans and other eukaryotes like *Candida* have a different and highly conserved functional squalene/phytoene synthase (ERG9) (41). Thus, the actual function of NDU1-like proteins in eukaryotic mitochondria is unknown. The human orthologue NDUFAF6 has been demonstrated to play an important role in the assembly of complex I through regulation of subunit ND1 biogenesis (39). Future work using docking and molecular dynamics approaches combined with biochemical assays will be needed to understand if the NDU1 cavity can accommodate and bind isoprenoid ligands. The identity of NDU1 substrates and its catalytic activity associated with complex I assembly, is yet to be determined. However, the fact that NDU1 is predicted to be membrane-localized could indicate that it interacts with amphiphilic or hydrophobic ligands, or possess a chaperone-like role in assembling integral membrane subunit proteins of complex I.

Despite overall similarities to human NDUFAF6, *C. albicans* NDU1 protein was found to be unique in its amino acid sequence. Compared to NDUFAF6, NDU1 is a significantly longer protein: 380 vs 333 amino acids (**Fig 5B**). The first long gap in NDU1 is part of the mitochondrial presequence; the human presequence is quite a bit longer than the *Candida* one. But the N-termini are removed upon import, so are not relevant to function. Note, however how several gaps have been introduced into the human sequence for the alignment to work because NDU1 is longer. The most compelling part of our study was that NDU1 protein sequence harbors three extra sets of amino acid inserts (**Fig 5C**), which are found only in CTG clade and very closely related fungi and missing from all other eukaryotes. One insert (insert 1) is positioned at the mouth of the cavity, while the other two inserts (insert 2 and 3) are structurally contiguous at the very bottom tip of the cavity (**Fig 6B**). Our gene deletion/complementation studies showed that insert 2, and 3, but not 1, are required for the function of NDU1. Because of the lack of insert 2, and 3 in the human NDUFAF6 orthologue, an overall low homology between the two proteins, and since NDU1 is required for virulence of *C. albicans*, the *C. albicans* NDU1 represents a highly desirable target for future novel therapeutic development to treat hematogenously disseminated candidiasis.

Inferring evolutionary mechanisms from genomic sequences with millions of years of divergence between them is inherently difficult. The idea that domain gains in eukaryotic proteins are directly mediated by gene duplication, followed by gene fusion and recombination may be the most plausible explanation. However, the *Candida* clade of species that are closely related to other Saccharomycotina did not undergo a whole genome duplication event (42). Interestingly, small-scale duplication events did occur in *C. albicans*, and genomic diversity continues to increase during exposure to stress (43–45), thereby facilitating functional diversification and providing greater phenotypic flexibility (46–48). Evolutionary diversity has also resulted in divergence in the post-transcriptional control of several processes in *Candida* spp. Specifically, mitochondrial protein synthesis and import is diverged in these fungi (12, 49, 50). *C. albicans*, which last shared a common ancestor with *S. cerevisiae* at least 300 million years ago contains a complete ETC, while *S. cerevisiae* is devoid of complex I (NADH: ubiquinone oxidoreductase) (51–53).

To conclude, our study reveals for the first time that following duplication, certain *C. albicans* genes may have acquired additional gene inserts to bolster protein-protein interactions. This unique evolutionary adaptation could also indicate lineage-specific changes in mitochondrial function, that likely are specific to how *Candida* adapts to nutritional stress. Why this selective acquisition of inserts is apparent only in NDU1 orthologues in the CTG clade fungi, and not in any other eukaryotes is a subject worth investigating. Certainly, sequences in the NDU1 protein that are different from its human orthologue can be harnessed as targets, for small molecule compounds that can dock to it and abrogate function. Severe virulence defects *in vivo* upon mitochondrial dysfunction due to NDU1 deletion suggest that inhibition of this target would be an effective way to combat fungal infections. Indeed, presence of orthologues of NDU1 in the multidrug resistant fungus *C. auris*, and in all other strains of *C. albicans* especially those resistant to antifungal drugs, means that mitochondrial inhibitors have a chance to act as pan-antifungal drugs.

## Methods

### Strains and culture conditions

Stock cultures of all strains were stored in 15% glycerol at −80°C. Strains were routinely grown under yeast conditions (media at 30°C) in YPG (1% yeast extract, 2% Bacto peptone, 2% glucose) or under filament-inducing conditions using RPMI medium (Sigma, St. Louis, MO) with MOPS (morpholinepropanesulfonic acid) buffer. For experiments requiring alternative sources for growth, YP was supplemented with either 2% of potassium acetate, sorbitol, glycerol or ethanol.

### Gene deletion and rescue of orf9.2500 (*NDU1*)

To generate orf19.2500 mutant, the orf19.2500 was replaced with the deletion cassette containing URA3 gene (54) as a selectable marker gene flanked with fragments corresponding to 500 bp upstream and downstream flanking sequences of the orf19.2500. We added KpnI and XhoI restriction sites to the ends of upstream fragment and NotI and SacII restrictions sites to the downstream fragment by PCR for cloning. The deletion cassette was released by KpnI and SacII restriction enzymes and transformed into BWP17 cells (55) followed by spreading the cells on uracil dropout medium. The heterozygous strain was confirmed by PCR and subjected to another round of gene deletion using a deletion cassette containing ARG4 as a selectable marker to prepare null mutant strain.

To compliment orf19.2500 mutant, we generated a complimented cassette containing a full length of orf19.2500, the nourseothricin resistance gene as a selectable marker and 500 bp of the terminator region of the orf19.2500 gene using pJK890 (55). The ORF19.2500 sequence was cloned with KpnI and ApaI restriction enzymes and the downstream part was cloned with NotI and SacII restriction enzymes into pJK890. The rescue cassette was released by KpnI and SacII restriction enzymes and transformed into the orf19.2500 mutant strain. The cells were spread on YPD containing 200ug/ml nourseothricin as a selection medium. The correct transformants were screened by PCR. To rescue the second allele of the ORF19.200 gene in this heterozygous strain, the nourseothricin resistance gene was looped out from the cells as described previously (6). Further, the cells were subjected to the rescue cassette again to receive the second allele of ORF19.2500 gene. The homozygous strain was confirmed by PCR.

### GFP and mCherry tagging combined with regulation of expression

We overexpressed orf19.2500 under Tetoff promoter and tagged the gene with GFP or mCherry at C-terminal. To do so, first the selective marker URA3 was replaced with ARG4 in pGS1245 (Tetoff-GFP-TetR-URA3) (56), then the full-length sequence of orf19.2500 was integrated into the plasmid via XhoI restriction enzyme. The plasmid was digested with AscI and transformed into orf19.2500-/- mutant to produce overexpressed strain. To tag the gene with mCherry, the GFP was replaced with mCherry sequence with XhoI and ClaI restriction enzymes. A similar approach was used also to overexpress the Tet-O promoter driven GFP-tagged, full length sequence of the human gene NDUFAF6 or the individual NDU1 insertion sequences in *C. albicans* orf19.2500 mutant strain.

### Growth rate determination

For cell dilutions spotted onto agar media as previously described (4), saturated overnight cultures were diluted in four to fivefold steps from an OD_600_ of 0.5. The stressors used were YPG agar plus 50 μg/ml calcofluor white, or 10 μg/ml congo red or 0.025% SDS. For growth curves in liquid media, saturated overnight cultures in YPD were washed once in 0.9% NaCl and diluted to an OD_600_ of 0.15 in 150 μL medium in flat-bottomed 96-well dishes. For growth assays OD_600_ readings were obtained every 60 min in a plate reader, and SDs of three technical replicates were calculated and graphed. For viability counts, *C. albicans* strains were inoculated at a concentration of 1×10^6^ cells/ml in 250 ml YPG medium. Every day up to 15 days, an aliquot of cells were recovered, diluted, counted using a hemocytometer, and plated on YPG agar plates. Colonies were counted, calculated and plotted, representing the viability of the cells over time.

### Biofilm growth and dispersal

Biofilms were grown both under static and flow conditions. For static growth, 1 ml of *C. albicans* cells (1 x 10^6^ cells/ml) was added to the wells of a 24-well microtiter plate and incubated overnight in RPMI, and the biofilms were gently washed two times (57). For enumeration of dispersed cells, static biofilm supernatants were collected after 24 h of growth, and turbidity (OD_600_) measured by a spectrophotometer. For growth under the flow system, biofilms were developed on silicone elastomer material, as previously described (58). At 24 h of biofilm growth, media flowing over the biofilms were collected, biofilm-dispersed cells present in the media counted using a hemocytometer, and plotted. Extent of attachment of the biofilm to its substrate was examined but gentle washing of the biofilm in the static model, or teasing the biofilm away from the substrate using a sterile needle. A small aliquot of the biofilm hyphae were also visualized under a phase contrast microscope (40X mag) to appreciate the extent of hyphae to lateral yeast growth.

### Assessment of phenotypic properties

#### Damage to HUVEC

Human Umbilical Cord Endothelial Cells (HUVEC) were isolated following an established protocol (3). The ability of *C. albicans* to damage human vascular endothelial cells was assessed by the CytoTox-96 assay (Promega, Madison, WI), which measures the release of lactate dehydrogenase (LDH) from dying cells. For these experiments, WT and mutant cells were diluted to various concentrations in HUVEC culture medium and were added to endothelial cells for 16 h incubation times at 37°C in the presence of 5% CO2. The amount of LDH released from the co-culture system was quantified by spectrophotometry. Uninfected cultures (control 1) and *C. albicans* alone (control 2) incubated under identical conditions were included as negative controls. The total amount of LDH released was estimated by treating control uninfected endothelial cells with 9% Triton X-100 for 1 hr. The LDH released in the presence of *C. albicans* was quantified by using the following formula: [(experimental – control 1 – control 2)/(total – control 1)] ×100. The values were expressed as percentages of the total amount of LDH released.

#### Cell membrane permeability

*Candida* strains were grown in YPG for 48 h and about 5 × 10^6^ cells were resuspended and washed twice in 1 ml of FDA buffer before supplementing with 50 nm FDA. A 200 μl volume of cell mixture with or without FDA was added to an optical-bottom 96-well plate. The kinetics of FDA uptake was recorded every 5 min for 30 reads with simultaneous shaking of samples in a plate reader with an excitation and emission wavelengths 485 and 535 nm, respectively. Data represent the fluorescence intensity over time.

#### Flow cytometry for cell component analysis

To stain mannan and chitin of the cell wall, *C*. *albicans* yeast cells grown for 48 h were washed in PBS and incubated in the dark with 25 μg/ml Concanavalin A to stain for α-mannopyranosyl or 5 μg/ml CFW for chitin for 30 min. The above stained cells were washed, fixed and differences intensity of the staining measured by flow cytometer at ~495/519 nm or 380/475 nm, respectively.

#### Transmission electron microscopy

*C. albicans* cells grown for 48 h were washed in PBS and then fixed in 4 ml fixative solution (3% paraformaldehyde, 2.5% glutaraldehyde, pH 7.2) for 24 h at 4°C. After post-fixation of samples with 1% phosphotungstic acid for 2 h, they were washed by distilled water, block-stained with uranyl acetate, dehydrated in alcohol, immersed in propylenoxide, and embedded in glycide-ether. Ultrathin sections were observed under a JEOL 100CX transmission electron microscope.

#### Fluconazole susceptibility

Fluconazole activity was assessed by Epsilometer test strips (Etest strips) (bioMérieux) according to the manufacturer’s instructions. A standardized cell suspension (a 0.5 McFarland standard) was used to create a confluent lawn across YPD agar plates prior to Etest strip placement, and the cells were then incubated at 30°C for 48 h.

##### Neutrophil killing

After obtaining institutional review board approved consent (The Lundquist protocol # 11672-07), neutrophils were isolated from blood collected from human volunteers using endotoxin-free Ficoll-Paque Plus reagent (Amersham Biosciences). The killing assay was carried out as described previously (59). Briefly, neutrophils were incubated with *C. albicans* yeast cells (neutrophil:fungus ratio, 5:1). Controls contained *C. albicans* without neutrophils. After 150 min, the mixtures were sonicated to disrupt neutrophils and the surviving fungi quantitatively cultured. The percentage of opsonophagocytic killing (OPK) was calculated by dividing the number of colony forming unit (CFU) in the tubes containing neutrophils by the number of CFU in tubes without neutrophils. *C. albicans* phagocytosis by neutrophils were visualized at 90 and 150 min, using a phase contrast microscope (40X mag).

##### Virulence assays

Animal studies were approved by the IACUC of The Lundquist Institute at Harbor–UCLA Medical Center, according to the NIH guidelines for animal housing and care. For the *C*. *albicans* infection *in vivo*, groups of CD1 female mice (6–8 weeks) were injected via lateral tail vein with 200 μl of a suspension containing indicated live *C*. *albicans* (2.5×10^5^ cells or 2.5×10^6^ cells) in sterile saline. Mice were monitored daily and differences in survival between infected groups were compared by the Log Rank test. Quantitative culturing of kidneys from mice infected with different strains of *Candida* was performed; mice were infected through tail veins, kidneys were harvested 2 and 5 days post infection, homogenized, serially diluted in 0.85% saline, and quantitatively cultured on YPG that contained 50 μg/ml chloramphenicol. Colonies were counted after incubation of the plates at 37°C for 24 to 48 hr, and results were expressed as log CFU per gram of infected organ.

Virulence assay under regulated gene expression conditions *in vivo*: Cultures of *C. albicans* strains for injection were grown overnight in YPD medium without doxycycline and incubated at 30°C. Cells (2.5 × 10^5^ cells in 200 μl of pyrogen-free saline solution per mouse) of the *C. albicans* tetO-NDU1/ndu1 strain were delivered by tail vein injection into two groups of mice, each consisting of eight 6-to-8-week-old female CD1 mice, with or without doxycycline in their drinking water (2 mg/ml in 5% sucrose). Cells of the control NDU1/NDU1 strain were injected at the same infecting dose into another group of animals (n = 8) with doxycycline in their drinking water. Pathogenicity of wild-type strains not containing any tetracycline-regulatable element is not affected by the presence or absence of doxycycline (60). Mice were monitored daily and differences in survival between infected groups were compared by the Log Rank test.

### Mitochondria associated assays

#### Sphaeroplast and mitochondria preparations

Cells were grown in 250 ml of YPD broth overnight at 30°C, washed once with cold water and once with buffer A (1 M sorbitol, 10 mM MgCl2, 50 mM Tris-HCl [pH 7.8]), centrifuged (5,000 rpm for 10 min). Cells were suspended in buffer A (50 ml) plus 30 mM dithiothreitol (DTT) for 15 min at 30°C with shaking (100 rpm) and then collected and suspended in buffer A with 1 mM DTT plus 100 mg of Zymolyase 20T (MP Biomedicals) per 10 g of pelleted cells. Shake cultures (100 rpm) were incubated at 30°C for 60 min or until 90% of cells were converted into spheroplasts (as determined by light microscopy). Spheroplasts were washed twice with buffer A. Crude preparations of mitochondria were isolated as previously described (32). Briefly, spheroplasts were suspended in 10 ml of cold buffer B (0.6 M mannitol, 1 mM EDTA, 0.5% bovine serum albumin [BSA], 1 mM phenylmethylsulfonyl fluoride [PMSF], 10 mM Tris-HCl [pH, 7.4]) and then broken mechanically using a Dounce homogenizer on an ice bath. Cell debris was removed by low-speed centrifugation (1,000 × g for 10 min). The supernatants containing mitochondria were centrifuged at 10,500 × g for 10 min, and the pellet was washed twice with 20 ml of ice-cold buffer C (0.6 M mannitol, 1 mM EDTA, 1% BSA, 10 mM Tris-HCl [pH 7.0]). Mitochondria were suspended in 1 ml of buffer D (0.6 M mannitol, 10 mM Tris-HCl, [pH 7.0]), and the protein content was determined by Bradford method.

#### Blue native PAGE

Mitochondrial protein was concentrated by vacuum centrifugation. Ten microliters of BN sample buffer (2X) was mixed with 20 μl of each sample (60 to 80 μg of protein) and loaded onto a BN-PAGE gradient gel (4 to 16%) (Invitrogen, Inc.). One ml of 2X BN sample buffer consisted of 1.5 M 6-aminohexanoic acid, 0.05 M bis-Tris (pH 7.0), 65 μl of 10% DMM, 20 μl of proteinase inhibitor mixture, and 100 μl of glycerol. Electrophoresis was performed in an X-Cell SureLock mini-cell system (Invitrogen) with 200 ml of cathode buffer in the upper (inner) buffer chamber and 150 ml of anode buffer in the lower (outer) buffer chamber. Electrophoresis was carried out at 4°C and 65 V for 1 h and then raised to 120 V overnight. An in-gel enzyme assay for CRC CI was accomplished as follows: gels were rinsed briefly twice with MilliQ water and equilibrated in 0.1 M Tris-HCl, pH 7.4 (reaction buffer), for 20 min. The gels were then incubated in fresh reaction buffer with 0.2 mM NADH–0.2% nitroblue tetrazolium (NBT) for 1 h. Reactions were stopped by fixing the gels in 45% methanol–10% (vol/vol) acetic acid, and then gels were destained overnight in the same solution. Image processing of gels was done using ImageJ software.

#### Enzymatic assay of CI

Mitochondrial protein was dissolved in 0.8 ml sterile water and incubated for 2 min at 37°C, then mixed with 0.2 ml of a solution containing 50 mM Tris pH 8.0, 5 mg /ml BSA, 0.24 mM KCN, 4 μM antimycin A and 0.8 mM NADH, the substrate for CI. The reaction was initiated by introducing an electron acceptor, 50 μM DB (2,3-dimethoxy-5-methyl-6-n-decyl-1,4 benzoquinone). Enzyme activity was followed by a decrease in absorbance of NADH at 340 nm minus that at 380 nm using an extinction coefficient of 5.5 mM^−1^cm^−1^

#### ROS measurement

Intracellular ROS production was detected by staining cells with 5 μM MitoSOX Red (Life Technologies) in DMSO. Cells from 25-ml cultures grown at 30°C overnight in YPD medium were collected and washed twice with PBS. The pellets were suspended to 1×10^6^ cells in 1 ml of PBS and treated with or without MitoSOX Red for 45 min at 30°C in the dark. Cell fluorescence in the presence of DMSO alone was used to verify that background fluorescence was similar per strain. Cells from each MitoSOX-treated sample were collected and washed twice with PBS after staining, and mean fluorescence for ROS was quantified.

#### Oxygen consumption rate (OCR) assay

OCR were measured under a Seahorse instrument (Seahorse Bioscience, MA) according to the manufacturer’s instructions. Isolated mitochondria from overnight grown WT, mutant and revertant cells were seeded into wells of a poly-d-lysine-coated XF96 spheroid plate containing 100 μL/well of warm assay medium (Seahorse XF base medium minimal DMEM, supplemented with 3 mM glucose and 0.1% FBS). 25 μl of mitochondrial suspension, containing three μg of protein for the succinate condition and pyruvate/malate condition, were added to a Seahorse 96-well plate and centrifuged (2000 g × 20 min × 4°C). After centrifugation, 155 μl assay buffer containing pyruvate (10 mM) in combination with malate (2 mM) or succinate (10 mM) and rotenone (2 μM) (all final concentrations and pH 7.2), were added, and the plate was analyzed at 37°C. Absolute OCR is presented as pmol O2 consumed/min/μg protein. Mitochondrial OCR was determined by subtracting the antimycin A (1μM, Sigma) and Rotenone (1 μM, Sigma)-sensitive OCR from the post-treatment OCR. Basal respiration was calculated in the presence of respiratory substrates (before ADP addition). Percentage inhibition was determined by dividing the post-treatment OCR with the basal mitochondrial OCR (antimycin A and Rotenone corrected) (61).

#### Mitochondrial membrane potential assay

The mitochondrial inner membrane potential (Δψm) was determined by staining with the membrane-permeable lipophilic cationic fluorochrome JC-1 (BD Biosciences, NJ). Overnight *C. albicans* cultures were washed, diluted to 1×10^6^ cells/ml of PBS, treated with JC-1 (3 μM final concentration) and incubated at 37°C for 30 min. Cells were washed and resuspended in 1 ml PBS and fluorescence dye accumulation measured using a flow cytometer equipped with a 488 nm argon excitation laser and 525 nm emission, and bandpass filters designed to detect green FITC dye (62).

---

**Figure S1**A. Growth curve of WT vs orf19.2500 mutant over 16 days

S1B. Colony size of mutant versus WT after 4 days of growth

S1C. Hyphal lengths of WT and mutant compared visually

S1D. Damage caused by WT and mutant cells to HUVEC cells measured after 24 h using the chromium release assay

**Figure S2**A. Estimation of cell wall mannan and chitin content by staining WT, mutant and revertant with ConA and calcofluor white, respectively, and measured by flow cytometry.

S2B. Visualization and measurement of the differences in thickness of the cell wall structure between WT and mutant cells, by transmission electron microscopy.

S2C. Quantitation of the extent of cell membrane permeability between WT and orf19.2500 mutant, in the presence and absence of fluconazole

S2D. Quantitation of the expression of ergosterol genes in WT and mutant cells

**Figure S3**A. Measurement of oxygen consumption rates of mitochondria isolated from WT, mutant and revertant strains, in presence of Complex II substrates succinate+rotenone

S3B. Determination on defect in mitochondrial membrane integrity in the WT, mutant and revertant strain, on growth in glucose or acetate

S3C. Survival of mice infected with a 10 fold higher infection dose of 2.5×10^6^ cells, of WT mutant and revertant cells.

**Figure S4**A. Surface display of 2 FPS bound in the large pocket in 5iys from *E. hirae*. Mg^++^ (green spheres), water molecules (red spheres)

S4B. Surface display of the pocket of c5iysA model while still showing FPS as they are positioned in 5iys. Note, while the pocket is in a different shape and the substrates cannot bind in the same orientations, the pocket is large enough to accommodate the 2 FPS.

S4C. Model predicted by Phyre2 shows c4hd1A (green), which is NDU1 modeled on 4hd1, a squalene synthase from *A. acidocaldarius*

S4D. Red highlighting of the identical residues between NDU1 and human NDUFAF6 (grey).

S4F. Structural model of the interaction of NDU1 with the surface of the membrane

**Figure S5**A. Phylogeny of NDU1. NDU1 belongs to the Trans_IPPS family. Proteins were aligned using TCOFFEE. Support values for nodes are from MrBayes (upper value) and RAxML (lower value). Putative orthologs of NDU1 form Cluster 1; PHYS (phytoene synthase) and SQS (squalene synthase) homologs form Cluster 2 and COQ1 (coenzyme Q1 synthase, decaprenyl diphosphate synthase) homologs form Cluster 3.

S5B. Flow cytometry data of *C. albicans* strains stained with MitoSox Red, an indicator of ROS activity. ROS production in NDU1 mutant overexpressing NDU1 without respective inserts were compared to ROS activity in WT and mutant strains. p<0.01 of the indicated conditions versus WT.

## Acknowledgements

We would like to thank Dr. Ana Traven, Professor, Monash University, Melbourne, Australia for her kind contribution on the preliminary assessment and proofreading of this manuscript.

## References

1. Nobile CJ, Johnson AD. Candida albicans Biofilms and Human Disease. Annu Rev Microbiol. 2015;69:71–92.

2. Uppuluri P, Lopez-Ribot JL. Go Forth and Colonize: Dispersal from Clinically Important Microbial Biofilms. PLoS Pathog. 2016;12(2):e1005397.

3. Uppuluri P, Chaturvedi AK, Srinivasan A, Banerjee M, Ramasubramaniam AK, Kohler JR, et al. Dispersion as an important step in the Candida albicans biofilm developmental cycle. PLoS Pathog. 2010;6(3):e1000828.

4. Uppuluri P, Acosta Zaldívar M, Anderson MZ, Dunn MJ, Berman J, Lopez Ribot JL, et al. Candida albicans Dispersed Cells Are Developmentally Distinct from Biofilm and Planktonic Cells. mBio. 2018;9(4):e01338–18.

5. Nobile CJ, Fox EP, Nett JE, Sorrells TR, Mitrovich QM, Hernday AD, et al. A recently evolved transcriptional network controls biofilm development in Candida albicans. Cell. 2012;148(1-2):126–38.

6. Shen J, Cowen LE, Griffin AM, Chan L, Kohler JR. The Candida albicans pescadillo homolog is required for normal hypha-to-yeast morphogenesis and yeast proliferation. Proc Natl Acad Sci U S A. 2008;105(52):20918–23.

7. Lemire BD. Evolution, structure and membrane association of NDUFAF6, an assembly factor for NADH:ubiquinone oxidoreductase (Complex I). Mitochondrion. 2017;35:13–22.

8. Ceri H, Olson ME, Stremick C, Read RR, Morck D, Buret A. The Calgary Biofilm Device: new technology for rapid determination of antibiotic susceptibilities of bacterial biofilms. J Clin Microbiol. 1999;37(6):1771–6.

9. Bahn YS, Staab J, Sundstrom P. Increased high-affinity phosphodiesterase PDE2 gene expression in germ tubes counteracts CAP1-dependent synthesis of cyclic AMP, limits hypha production and promotes virulence of Candida albicans. Mol Microbiol. 2003;50(2):391–409.

10. Day M. Yeast petites and small colony variants: for everything there is a season. Adv Appl Microbiol. 2013;85:1–41.

11. Rodrigues ML. The Multifunctional Fungal Ergosterol. mBio. 2018;9(5):e01755–18.

12. Sun N, Parrish RS, Calderone RA, Fonzi WA. Unique, Diverged, and Conserved Mitochondrial Functions Influencing <span class=“named-content genus-species” id=“named-content-1”>Candida albicans</span> Respiration. mBio. 2019;10(3):e00300–19.

13. Zhao R-Z, Jiang S, Zhang L, Yu Z-B. Mitochondrial electron transport chain, ROS generation and uncoupling (Review). Int J Mol Med. 2019;44(1):3–15.

14. Lorenz MC, Fink GR. The glyoxylate cycle is required for fungal virulence. Nature. 2001;412(6842):83–6.

15. Claros MG, Vincens P. Computational method to predict mitochondrially imported proteins and their targeting sequences. Eur J Biochem. 1996;241(3):779–86.

16. Kelley LA, Mezulis S, Yates CM, Wass MN, Sternberg MJ. The Phyre2 web portal for protein modeling, prediction and analysis. Nat Protoc. 2015;10(6):845–58.

17. Dundas J, Ouyang Z, Tseng J, Binkowski A, Turpaz Y, Liang J. CASTp: computed atlas of surface topography of proteins with structural and topographical mapping of functionally annotated residues. Nucleic Acids Res. 2006;34(Web Server issue):W116–8.

18. Yang J, Yan R, Roy A, Xu D, Poisson J, Zhang Y. The I-TASSER Suite: protein structure and function prediction. Nat Methods. 2015;12(1):7–8.

19. Katoh K, Standley DM. MAFFT multiple sequence alignment software version 7: improvements in performance and usability. Mol Biol Evol. 2013;30(4):772–80.

20. Nguyen LT, Schmidt HA, von Haeseler A, Minh BQ. IQ-TREE: a fast and effective stochastic algorithm for estimating maximum-likelihood phylogenies. Mol Biol Evol. 2015;32(1):268–74.

21. Lomize MA, Pogozheva ID, Joo H, Mosberg HI, Lomize AL. OPM database and PPM web server: resources for positioning of proteins in membranes. Nucleic Acids Res. 2012;40(Database issue):D370–6.

22. Ramírez MA, Lorenz MC. Mutations in alternative carbon utilization pathways in Candida albicans attenuate virulence and confer pleiotropic phenotypes. Eukaryotic cell. 2007;6(2):280–90.

23. Hatab MA, Whittaker PA. Isolation and characterization of respiration-deficient mutants from the pathogenic yeast Candida albicans. Antonie Van Leeuwenhoek. 1992;61(3):207–19.

24. Calderone R, Li D, Traven A. System-level impact of mitochondria on fungal virulence: to metabolism and beyond. FEMS Yeast Res. 2015;15(4):fov027.

25. She X, Calderone R, Kruppa M, Lowman D, Williams D, Zhang L, et al. Cell Wall N-Linked Mannoprotein Biosynthesis Requires Goa1p, a Putative Regulator of Mitochondrial Complex I in Candida albicans. PLoS One. 2016;11(1):e0147175.

26. She X, Zhang L, Chen H, Calderone R, Li D. Cell surface changes in the Candida albicans mitochondrial mutant goa1Delta are associated with reduced recognition by innate immune cells. Cell Microbiol. 2013;15(9):1572–84.

27. Qu Y, Jelicic B, Pettolino F, Perry A, Lo TL, Hewitt VL, et al. Mitochondrial sorting and assembly machinery subunit Sam37 in Candida albicans: insight into the roles of mitochondria in fitness, cell wall integrity, and virulence. Eukaryot Cell. 2012;11(4):532–44.

28. Fox EP, Cowley ES, Nobile CJ, Hartooni N, Newman DK, Johnson AD. Anaerobic bacteria grow within Candida albicans biofilms and induce biofilm formation in suspension cultures. Current biology : CB. 2014;24(20):2411–6.

29. Morales DK, Grahl N, Okegbe C, Dietrich LEP, Jacobs NJ, Hogan DA. Control of *Candida albicans* Metabolism and Biofilm Formation by *Pseudomonas aeruginosa* Phenazines. mBio. 2013;4(1):e00526–12.

30. Askew C, Sellam A, Epp E, Hogues H, Mullick A, Nantel A, et al. Transcriptional regulation of carbohydrate metabolism in the human pathogen Candida albicans. PLoS Pathog. 2009;5(10):e1000612.

31. Rodaki A, Bohovych IM, Enjalbert B, Young T, Odds FC, Gow NA, et al. Glucose promotes stress resistance in the fungal pathogen Candida albicans. Mol Biol Cell. 2009;20(22):4845–55.

32. Li D, Chen H, Florentino A, Alex D, Sikorski P, Fonzi WA, et al. Enzymatic dysfunction of mitochondrial complex I of the Candida albicans goa1 mutant is associated with increased reactive oxidants and cell death. Eukaryot Cell. 2011;10(5):672–82.

33. Vincent BM, Langlois J-B, Srinivas R, Lancaster AK, Scherz-Shouval R, Whitesell L, et al. A Fungal-Selective Cytochrome bc(1) Inhibitor Impairs Virulence and Prevents the Evolution of Drug Resistance. Cell Chem Biol. 2016;23(8):978–91.

34. Li S-X, Song Y-J, Zhang Y-S, Wu H-T, Guo H, Zhu K-J, et al. Mitochondrial Complex V α Subunit Is Critical for Candida albicans Pathogenicity through Modulating Multiple Virulence Properties. Front Microbiol. 20178:285-.

35. Bambach A, Fernandes MP, Ghosh A, Kruppa M, Alex D, Li D, et al. Goa1p of Candida albicans localizes to the mitochondria during stress and is required for mitochondrial function and virulence. Eukaryot Cell. 2009;8(11):1706–20.

36. Li D, She X, Calderone R. Functional diversity of complex I subunits in Candida albicans mitochondria. Curr Genet. 2016;62(1):87–95.

37. Santos MA, Gomes AC, Santos MC, Carreto LC, Moura GR. The genetic code of the fungal CTG clade. Comptes rendus biologies. 2011;334(8–9):607–11.

38. Pandit J, Danley DE, Schulte GK, Mazzalupo S, Pauly TA, Hayward CM, et al. Crystal structure of human squalene synthase. A key enzyme in cholesterol biosynthesis. J Biol Chem. 2000;275(39):30610–7.

39. McKenzie M, Tucker EJ, Compton AG, Lazarou M, George C, Thorburn DR, et al. Mutations in the gene encoding C8orf38 block complex I assembly by inhibiting production of the mitochondria-encoded subunit ND1. Journal of molecular biology. 2011;414(3):413–26.

40. Liu CI, Jeng WY, Chang WJ, Shih MF, Ko TP, Wang AH. Structural insights into the catalytic mechanism of human squalene synthase. Acta Crystallogr D Biol Crystallogr. 2014;70(Pt 2):231–41.

41. Nakayama H, Izuta M, Nakayama N, Arisawa M, Aoki Y. Depletion of the squalene synthase (ERG9) gene does not impair growth of Candida glabrata in mice. Antimicrob Agents Chemother. 2000;44(9):2411–8.

42. Rozpędowska E, Galafassi S, Johansson L, Hagman A, Piškur J, Compagno C. Candida albicans--a pre-whole genome duplication yeast--is predominantly aerobic and a poor ethanol producer. FEMS Yeast Res. 2011;11(3):285–91.

43. Bouchonville K, Forche A, Tang KE, Selmecki A, Berman J. Aneuploid chromosomes are highly unstable during DNA transformation of Candida albicans. Eukaryot Cell. 2009;8(10):1554–66.

44. Forche A, Alby K, Schaefer D, Johnson AD, Berman J, Bennett RJ. The parasexual cycle in Candida albicans provides an alternative pathway to meiosis for the formation of recombinant strains. PLoS Biol. 2008;6(5):e110.

45. Forche A, Abbey D, Pisithkul T, Weinzierl MA, Ringstrom T, Bruck D, et al. Stress alters rates and types of loss of heterozygosity in Candida albicans. mBio. 2011;2(4):e00129–11.

46. Anderson MZ, Baller JA, Dulmage K, Wigen L, Berman J. The three clades of the telomere-associated TLO gene family of Candida albicans have different splicing, localization, and expression features. Eukaryotic cell. 2012;11(10):1268–75.

47. Butler G, Rasmussen MD, Lin MF, Santos MAS, Sakthikumar S, Munro CA, et al. Evolution of pathogenicity and sexual reproduction in eight Candida genomes. Nature. 2009;459(7247):657–62.

48. Dunn MJ, Kinney GM, Washington PM, Berman J, Anderson MZ. unctional diversification accompanies gene family expansion of MED2 homologs in Candida albicans. PLoS genetics. 2018;14(4):e1007326–e.

49. Dagley MJ, Gentle IE, Beilharz TH, Pettolino FA, Djordjevic JT, Lo TL, et al. Cell wall integrity is linked to mitochondria and phospholipid homeostasis in Candida albicans through the activity of the post-transcriptional regulator Ccr4-Pop2. Mol Microbiol. 2011;79(4):968–89.

50. Hewitt VL, Heinz E, Shingu-Vazquez M, Qu Y, Jelicic B, Lo TL, et al. A model system for mitochondrial biogenesis reveals evolutionary rewiring of protein import and membrane assembly pathways. Proc Natl Acad Sci U S A. 2012;109(49):E3358–66.

51. Gabaldón T, Rainey D, Huynen MA. Tracing the evolution of a large protein complex in the eukaryotes, NADH:ubiquinone oxidoreductase (Complex I). Journal of molecular biology. 2005;348(4):857–70.

52. Marcet-Houben M, Marceddu G, Gabaldón T. Phylogenomics of the oxidative phosphorylation in fungi reveals extensive gene duplication followed by functional divergence. BMC evolutionary biology. 2009;9:295.

53. Lavín JL, Oguiza JA, Ramírez L, Pisabarro AG. Comparative genomics of the oxidative phosphorylation system in fungi. Fungal genetics and biology : FG & B. 2008;45(9):1248–56.

54. Mamouei Z, Zeng G, Wang YM, Wang Y. Candida albicans possess a highly versatile and dynamic high-affinity iron transport system important for its commensal-pathogenic lifestyle. Mol Microbiol. 2017;106(6):986–98.

55. Wilson RB, Davis D, Mitchell AP. Rapid hypothesis testing with Candida albicans through gene disruption with short homology regions. J Bacteriol. 1999;181(6):1868–74.

56. Zheng XD, Wang YM, Wang Y. CaSPA2 is important for polarity establishment and maintenance in Candida albicans. Mol Microbiol. 2003;49(5):1391–405.

57. Pierce CG, Uppuluri P, Tristan AR, Wormley FL, Jr., Mowat E, Ramage G, et al. A simple and reproducible 96-well plate-based method for the formation of fungal biofilms and its application to antifungal susceptibility testing. Nat Protoc. 2008;3(9):1494–500.

58. Uppuluri P, Lopez-Ribot JL. An easy and economical in vitro method for the formation of Candida albicans biofilms under continuous conditions of flow. Virulence. 2010;1(6):483–7.

59. Uppuluri P, Singh S, Alqarihi A, Schmidt CS, Hennessey JP, Jr., Yeaman MR, et al. Human Anti-Als3p Antibodies Are Surrogate Markers of NDV-3A Vaccine Efficacy Against Recurrent Vulvovaginal Candidiasis. Front Immunol. 2018;9:1349.

60. Chaturvedi AK, Lazzell AL, Saville SP, Wormley FL, Jr., Monteagudo C, Lopez-Ribot JL. Validation of the tetracycline regulatable gene expression system for the study of the pathogenesis of infectious disease. PLoS One. 2011;6(5):e20449.

61. Andersen JV, Jakobsen E, Waagepetersen HS, Aldana BI. Distinct differences in rates of oxygen consumption and ATP synthesis of regionally isolated non-synaptic mouse brain mitochondria. J Neurosci Res. 2019;97(8):961–74.

62. Sivandzade F, Bhalerao A, Cucullo L. Analysis of the Mitochondrial Membrane Potential Using the Cationic JC-1 Dye as a Sensitive Fluorescent Probe. Bio protoc. 2019;9(1).

